# Exploring Arylidene-Indolinone Ligands of Autophagy Proteins LC3B and GABARAP

**DOI:** 10.1101/2024.02.25.581879

**Authors:** Alexandria N. Leveille, Thomas Schwarzrock, Hawley Brown, Bennett True, Joanet Plasencia, Philipp Neudecker, Alina Üffing, Oliver H. Weiergräber, Dieter Willbold, Joshua A. Kritzer

## Abstract

We report the first structure-activity studies of arylidene-indolinone compound **GW5074** which was reported as a ligand of autophagy-related protein LC3B. The literature has conflicting information on the binding affinity of this compound and there is some debate regarding its use as a component of autophagy-dependent degrader compounds. We developed an AlphaScreen assay to measure competitive inhibition of the binding of known peptide ligands to LC3B and its paralog GABARAP. 18 analogs were synthesized and tested against both proteins. Inhibitory potencies were found to be in the mid- to high micromolar range. 2D-NMR data revealed the binding site on GABARAP as hydrophobic pocket 1, where native peptide ligands bind with an aromatic side chain. Our results suggest that **GW5074** binds LC3B and GABARAP with micromolar affinity. These affinities could support further exploration in targeted protein degradation, but only if off-target effects and poor solubility can be appropriately addressed.

---

Macroautophagy is a process by which intracellular cargo, including proteins, protein aggregates, and organelles, is recruited to and degraded by the lysosome. Macroautophagy (referred to here as simply “autophagy”) is essential for maintaining cellular homeostasis. In advanced cancers, autophagy contributes to disease progression through multiple mechanisms including immune evasion,^1,2^ metabolic adaptation,^3^ and accelerated metastasis.^4,5^ Additionally, autophagy is typically upregulated in response to DNA-damaging agents, and genetic studies have shown that inhibiting autophagy re-sensitizes late-stage cancers to cisplatin treatment.^6–8^ Therefore, autophagy inhibitors are a promising area for novel combination therapies.^9^ Unfortunately, commonly used autophagy inhibitors such as hydroxychloroquine are not specific to autophagy, they have many side effects, and they can have dose-limiting toxicity.^10,11^ Targeting protein-protein interactions involved in autophagy is a good strategy for developing selective inhibitors.^9^ Specifically, LC3/GABARAP family proteins mediate protein-protein interactions at every step of the autophagy pathway.^12^ Genetic knockdowns and knockouts of LC3/GABARAP proteins inhibit autophagy selectively, demonstrating that this family of autophagy proteins is a promising drug target.^13–16^ At high concentrations, ligands for LC3/GABARAP proteins should be effective autophagy inhibitors, but at lower concentrations they could also be used as components of targeted degrader compounds. Such degrader compounds have been termed autophagy-targeting chimeras (AUTACs) or autophagosome-tethering chimeras (ATTECs), among other terms. These compounds tether proteins or cellular components of interest to the autophagosome, leading to the degradation of the liganded protein, organelle, or cellular component.^17–29^

**GW5074** (Fig. 1, hereafter called compound **1**) was one of the earliest small molecules reported to bind LC3B, the most well-studied LC3/GABARAP protein. It was discovered in 2019 by Li and coworkers as a compound that induced degradation of aggregated 72Q-huntingtin.^19^ Since that report, **1** has been used for several applications. For example, in 2021, Li and coworkers attached **1** to a lipid droplet-binding compound to produce a chimeric compound that degraded lipid droplets.^20^ Also in 2021, Fu et al. attached **1** to the BRD4 ligand JQ1, producing a chimeric compound that degraded BRD4.^17^ More recently, Gu and coworkers reported an ATTEC that incorporated **1** to degrade the protein PCSK9 as a potential atherosclerosis therapy.^22^ Similarly, Dong and coworkers recently described an ATTEC using **1** to degrade PDEδ, an emerging target for pancreatic cancer therapy.^29^

**Figure 1.**
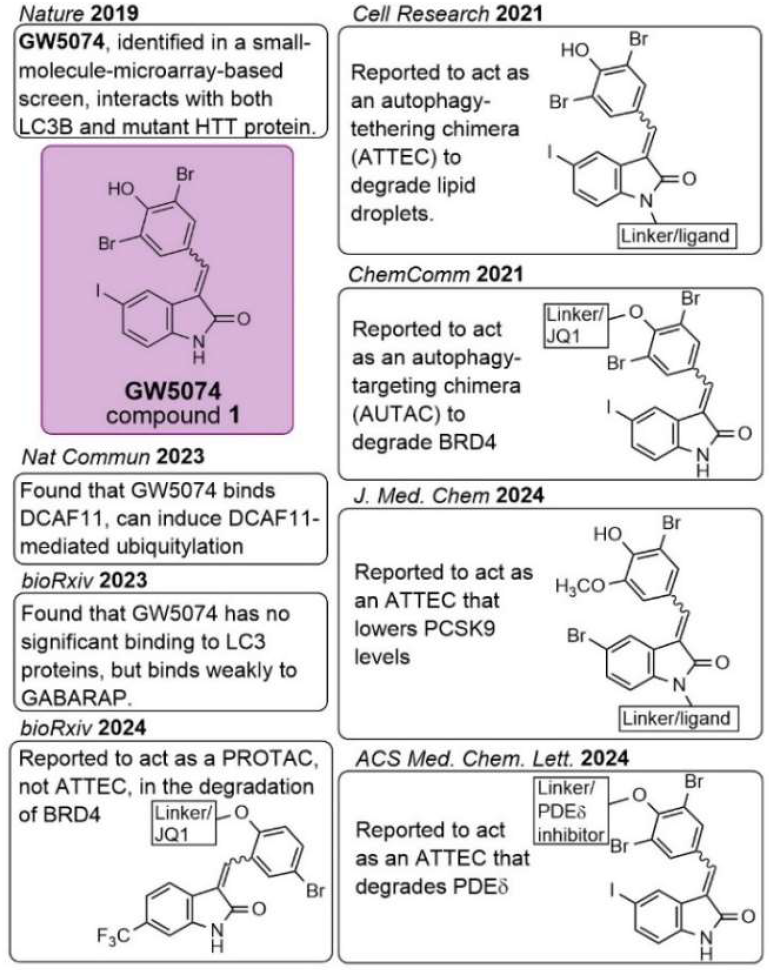
Summary of findings related to compound 1, also known as GW5074, in prior literature. Most arylidene-indolinones analogous to compound **1** are interconverting diastereomers with respect to the double bond in the arylidene.

These findings suggested that **1** could be used in selective and modular fashion to bind LC3/GABARAP proteins and induce the degradation of other proteins and cellular components via autophagy. However, other work has questioned the mechanism of degradation for ATTECs incorporating **1**.

Recent findings by Winter, Waldmann, and coworkers found that **1** is a selective, covalent ligand of the E3 ligase DCAF11.^30^ They further showed that its ability to direct targeted protein degradation is dependent on the proteasome and not lysosomal function, consistent with DCAF11 engagement in cells. Similarly, a recent preprint by Hong, Wang, Tian, Li, and coworkers found that a heterobifunctional molecule of JQ1 and a **1** derivative degraded BRD4 through recruitment of the E3 complex CRL4 which contains DCAF11.^31^ Thus, degradation by chimeric compounds incorporating **1** may not occur via an autophagy-dependent mechanism. Notably, while the report from Winter, Waldmann, and coworkers tested many analogs of **1** for DCAF11 engagement, none were tested for LC3/GABARAP binding or inhibition.

Compound **1** was derived from a screen of known bioactive compounds and it has not undergone any reported optimization or structure-activity relationship studies for LC3/GABARAP inhibition. This lack of information contributes to the questions surrounding the ability of **1** to inhibit LC3/GABARAP proteins. Even the published data on **1** binding to LC3/GABARAP proteins have been contradictory (Fig. 1). The binding affinity of **1** for recombinant LC3B was reported in different papers as 0.468 µM (measured by small molecule microarray with a scanning oblique-incidence reflectivity difference microscope),^19^ 8.9 µM (measured by surface plasmon resonance),^17^ and greater than 200 µM (measured by 2D-NMR titration).^32^ Overall, these contradictory findings called into question the actual binding affinity of **1** for LC3B and other family members. A recent report by Knapp, Rogov, and coworkers more directly addressed this question using competition fluorescence polarization, NMR titration, and NanoBRET assays. They found that **1** binds weakly to GABARAPL2 and has weak, if any, binding to LC3B.^32^ In that work, the authors suggested that these activities were too weak to account for the compound’s ability to mediate targeted degradation.

Concurrently with these more recent studies, we took up the question of whether **1** binds recombinant LC3B and GABARAP and whether it inhibits their interactions with representative ligands derived from native binding partners. We also sought to uncover structure-activity relationships for these inhibitory activities. We began by developing more reliable inhibition assays for recombinantly expressed LC3B and GABARAP. We first tried to test **1** in competitive fluorescence polarization assays that were previously developed in the Kritzer lab.^33^ However, the compounds had background fluorescence which interfered with the assay. As a convenient alternative to biolayer interferometry assays developed by us and others,^33–35^ we developed a solution-phase competition assay using AlphaScreen. The AlphaScreen assay (Fig. 2) reports on the inhibitor’s ability to block binding of known peptide ligands of LC3B or GABARAP which bind in the canonical protein-protein interaction site responsible for these proteins’ functions. For GABARAP, the ligand was biotin-labeled K1 peptide, which has a K_d_ of 55 ± 8 nM as measured by biolayer interferometry.^34,36^ For LC3B, the ligand was a modified, biotin-labeled FYCO1 peptide, FYCO1S, which has a K_d_ of 330 ± 30 nM as measured by biolayer interferometry.^33^ Using untagged versions of these tracer peptides as positive controls, we measured dose dependent inhibition in AlphaScreen with IC_50_ values of 55 ± 3 nM for K1 inhibiting GABARAP’s interaction with biotinylated K1, and 56 ± 4 nM for FYCO1S inhibiting LC3B’s interaction with biotinylated FYCO1S. These controls demonstrate that this assay measures the dose-dependent inhibition of the protein-peptide interactions relevant for LC3/GABARAP functions in autophagy. These new assays are similar to an AlphaScreen assay reported in 2021 by Proschak and coworkers, who measured the disruption of binding between GST-LC3B and biotin-LIRtide, a short peptide derived from LC3-Interacting Region (LIR) of p62.^37^

**Figure 2.**
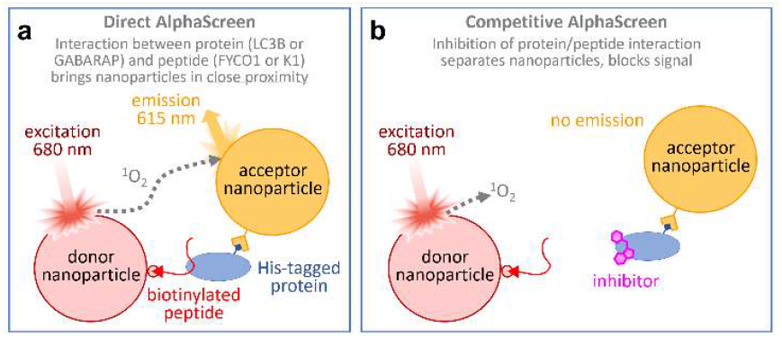
Newly developed AlphaScreen assays. One assay was used to measure inhibitory potency for inhibition of LC3B binding to the dye-labeled peptide ligand FYCO1S, and a similar assay was used to measure inhibitory potency for inhibition of GABARAP binding to the dye-labeled peptide ligand K1. Peptide ligands were immobilized on streptavidin-functionalized donor beads and recombinant LC3B or GABARAP was immobilized on Ni-NTA-functionalized acceptor beads. Additional assay details provided in Supporting Information.

Using the AlphaScreen assays, we measured the IC_50_ of **1** at 9.1 µM for GABARAP and 4.4 µM for LC3B. We also noted that **1** was only soluble to roughly 140 μM in the aqueous buffer used for AlphaScreen. We surmised that prior irreproducibility in determining binding affinities of **1** could have been due to insolubility in aqueous solution. Because **1** had substantial absorbance of visible light (λ_max_ at 498 nm), we calculated an extinction coefficient for **1** in DMSO and subsequently verified the concentration of **1** every time it was dissolved in aqueous solution by lyophilizing the aqueous working stock, resolubilizing in DMSO, and checking absorbance. This process was used for every analog of **1** to ensure accurate concentrations and avoid misinterpretations due to insolubility (Table S1).

We next developed structure-activity relationships by altering one functional group at a time on **1** (Fig. 3). Commercially available oxindoles and aldehydes readily underwent an aldol condensation in the presence of a base and ethanol, allowing us to produce a library of 18 analogs in total. Compounds described are a mixture of E/Z stereoisomers. Though a single stereoisomer was typically isolated following the aldol condensation, we observed that racemization readily occurred at room temperature (Fig. S2). We first prepared analogs with changes to the arylidene portion of **1** (Fig. 3). We observed that removal of all functional groups from the arylidene (**2a**) resulted in loss of inhibition for both proteins. Removing the bromo groups at C10 and C12 while retaining the hydroxyl group at C11 (**2b**) decreased solubility and resulted in loss of inhibition for both proteins when tested up to 10 μM. Removal of the hydroxyl at C11 while retaining the bromo groups at C10 and C12 (**2c**) led to a compound that was too insoluble to allow measurement of IC_50_ values. Moving the hydroxy group to the C9/C13 position (**2d**) resulted in decreased (compared to compound **1**) but measurable inhibition for both proteins. After methylation of the hydroxyl group at the C4 position (**2e**) solubility was insufficient for inhibition measurements.

**Figure 3.**
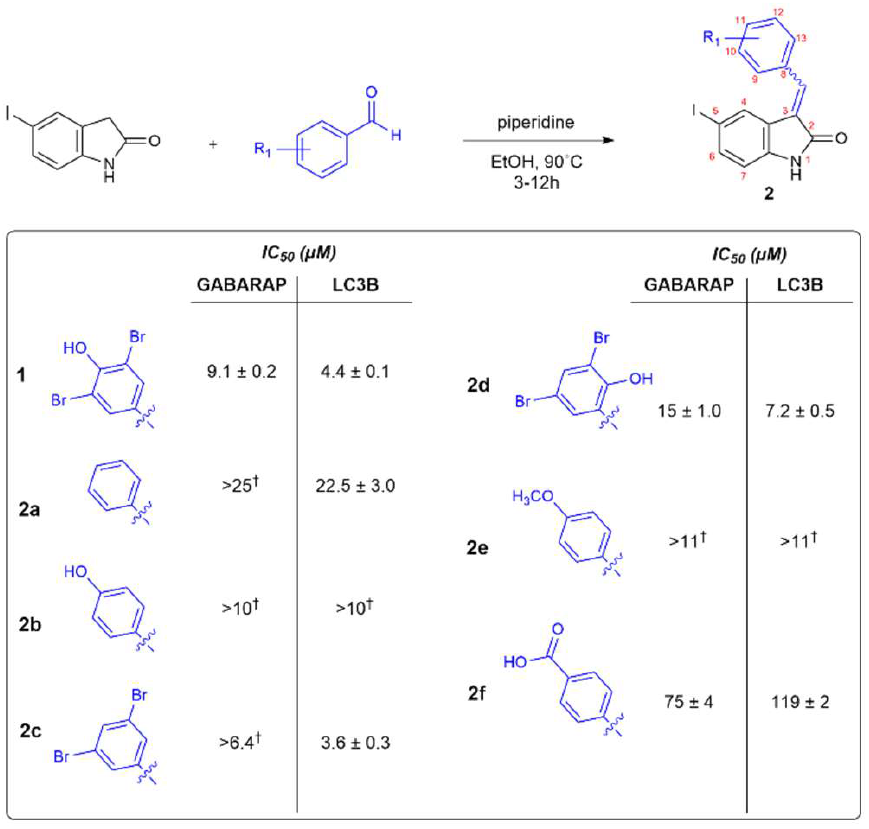
Exploring deletion and movement of functional groups on the arylidene portion of compound 1. Complete compound characterization and assay data are provided in Supporting Information. ^†^No inhibition observed up to the highest concentration tested, which was 60% of the solubility limit; solubility limits shown in Table S1.

When the *para*-hydroxyl group was replaced with a carboxylic acid (**2f**), binding for each protein was measurable though drastically decreased compared to **1**. Extinction coefficients, λ_max_ values, and solubility of each compound in aqueous solution can be found in Table S1.

We then explored substitutions on the oxindole portion of **1**. Removal of the iodine from the oxindole (**2g**) resulted in significantly decreased binding to both proteins compared to **1** (Fig. 4). Movement of the iodine from the C5 to the C6 position of the oxindole (**2h**) resulted in comparable binding as **1** to LC3B and slightly decreased binding to GABARAP. Substitution of iodine with a bromine at C5 (**2i**) resulted in slightly decreased binding to both proteins. Replacement of an electron-withdrawing halogen with an electron-donating methyl group at C6 resulted in decreased inhibition for both proteins (**2j**). Placement of methyl acetate at C6 (**2k**) resulted in increased inhibition of GABARAP though a slight decrease in inhibition of LC3B. The placement of a chlorine at both C5 and C6 resulted in measurable, though decreased, binding for LC3B but no measurable data for GABARAP (**2l**).

**Figure 4.**
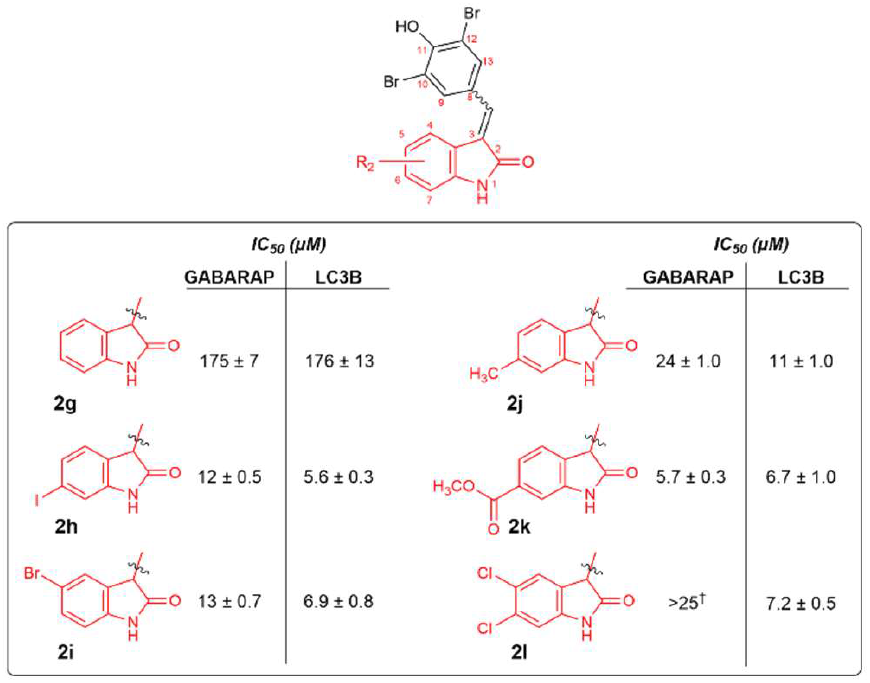
Exploring deletion and movement of functional groups on the oxindole portion of compound 1. Complete compound characterization and assay data are provided in Supporting Information. ^†^No inhibition observed up to the highest concentration tested, which was 60% of the solubility limit; solubility limits shown in Table S1.

After acquiring these initial structure-activity relationships, we next sought to alter the scaffold of **1** starting with substituting the arylidene with various commercially available heterocycles (Fig. 5). Unfortunately, some arylidene substitutions resulted in poor solubility with starting concentrations that were too low to measure any detectable inhibition for either protein. These substitutions included para-trifluoromethyl biphenyl (**2m**), thiazole (**2n**), and pyrrole (**2p**). 4-methyl thiazole (**2o**), imidazole (**2q**), and imidazo[1,2]pyridine (**2r**) substitutions were more soluble (soluble to 19, 118, and 22 μM, respectively). **2r** had no measurable inhibition for either protein up to 60% of its solubility limit, while **2o** and **2q** had some inhibition for LC3B but not GABARAP at the highest concentrations tested.

**Figure 5.**
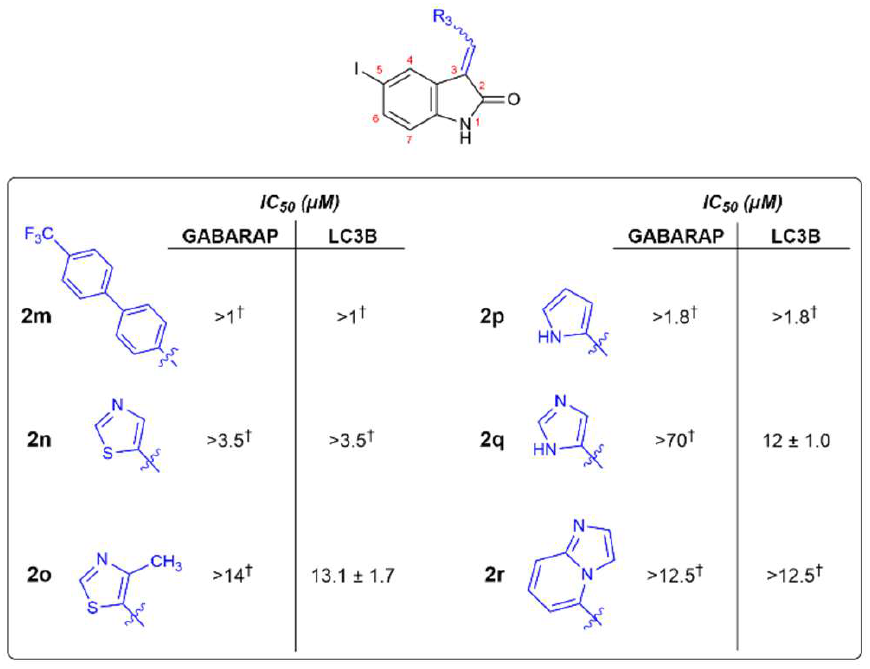
Exploring substitutions of heterocycles within the arylidene group.

The structure-activity relationships of **1** with respect to solubility and GABARAP inhibition are summarized in Fig. 6. The selectivity determinants of peptide and protein binding to LC3B and GABARAP have been extensively explored,^33–35,38–41^ but few studies have explored selectivity of small molecule binding or inhibition.^32,37,42^ Compound **1** has roughly 2-fold greater inhibitory potency for LC3B compared to GABARAP, and this selectivity was shared by nearly all analogs. Notably, compound **2k** which replaced the iodo group at C5 with a methyl acetate group at C6 had similar overall potency to compound **1** but had roughly equal inhibitory potencies for both LC3B and GABARAP.

**Figure 6.**
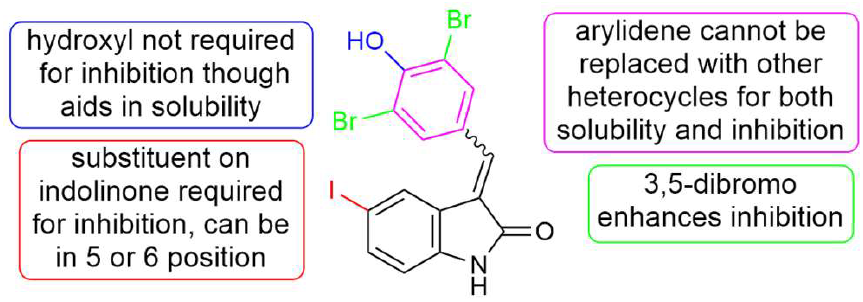
Summary of structure-activity relationships of **1**.

To rule out an entirely nonspecific mode of action, we also tested **1** in an assay for inhibition of beta-lactamase activity. If the mode of inhibition for LC3/GABARAP proteins was nonspecific, for instance via colloidal aggregation of the compounds leading to protein denaturation,^43,44^ we would expect to see inhibition of an unrelated protein at similar concentrations. We observed no effect on beta-lactamase activity for compound **1** at concentrations up to 25 µM (Fig. S1).

The binding site of compound **1** on GABARAP was investigated in more detail by titrating [U-^15^N] GABARAP with increasing amounts of **1** and monitoring GABARAP using 2D [^1^H,^15^N] HSQC NMR (Fig. 7). Ligand binding led to sizeable chemical shift perturbations (CSPs), several of which (E8, E17, E19, R22, K23, Y25, V29, V31, I32, V33, K48, Y49, L50, V51, S53, H70, F103, F104, Y106) systematically exceeded twice the root-mean-square CSP (Fig. 7a). Y49 was affected by strong line broadening in addition to a particularly large CSP. Mapping the chemical shift perturbations onto the molecular surface of GABARAP clearly shows that **1** binds to the N-terminal hydrophobic pocket of the two-pocket LIR binding site (Fig. 7c). This hydrophobic pocket, commonly referred to as HP1, is where aromatic residues from peptide ligands bind. This finding makes sense since the scaffold of **1** is made up of two aromatic ring systems and inhibitory potency was especially sensitive to replacements on the arylidene. Given that residues lining HP1 are highly conserved across LC3/GABARAP subfamily members,^40^ the binding of **1** in this pocket is consistent with the mild selectivity found for the inhibitory compounds we tested.

**Figure 7.**
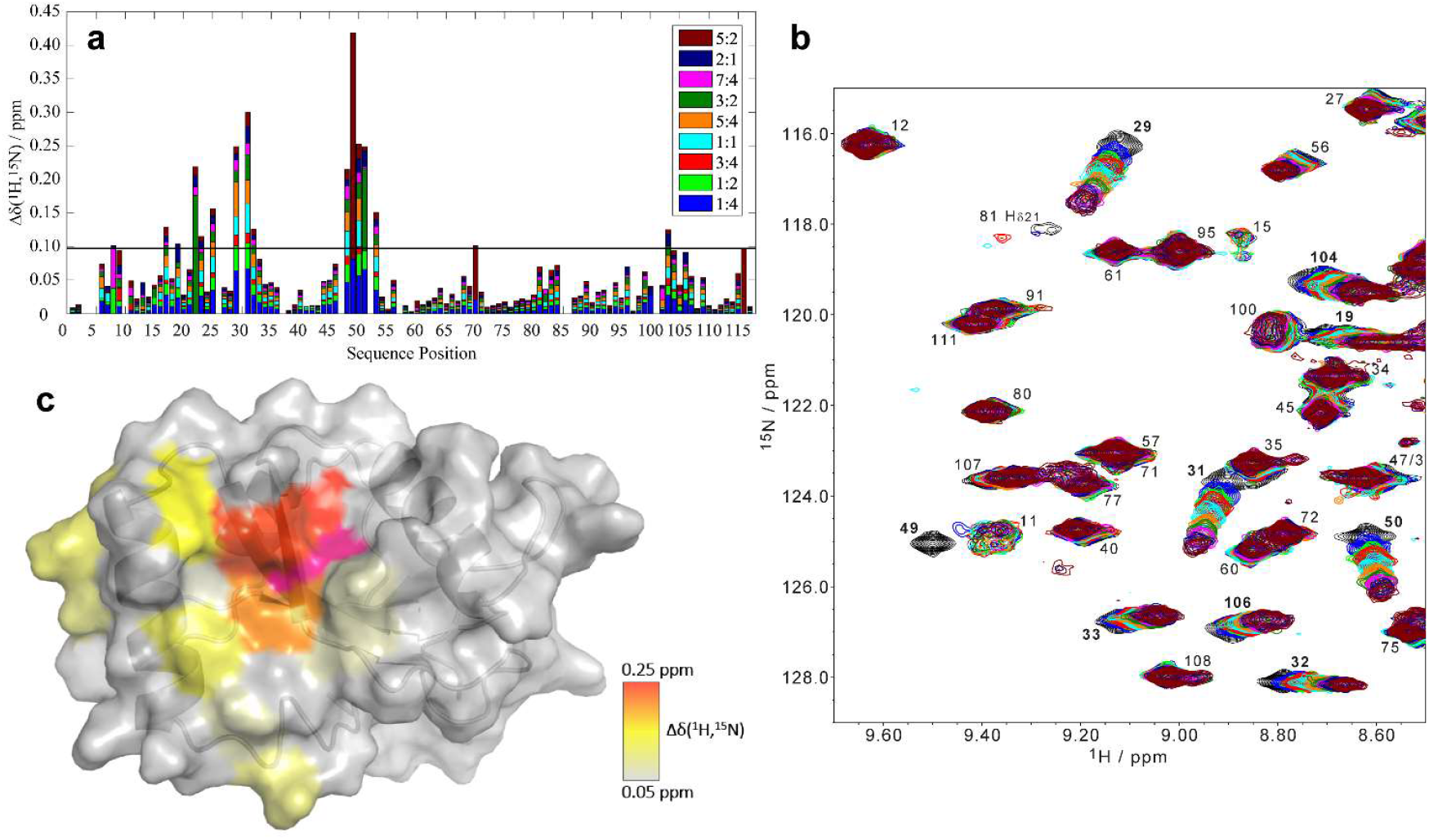
Titration of GABARAP with compound **1** as monitored by NMR spectroscopy. a) Chemical shift perturbations (CSPs) for backbone amide resonances of GABARAP incubated with **1** at the indicated stoichiometric ratios (**1**:GABARAP). The horizontal line indicates twice the root-mean-square CSP variation (0.0972 ppm) at 5:2 excess of **1**.b) Section of the [^1^H,^15^N] HSQC showing chemical shift perturbations with increasing concentrations of **1**. Spectra are color-coded to match the ratios shown in panel a, and the spectrum for GABARAP in the absence of **1** is shown in black. Full spectra are provided in the Supplementary Information. c) Molecular surface of GABARAP (PDB 1KOT)^48^ with residues colored according to their CSPs observed at a 5:2 (**1**:GABARAP) ratio. Y49, highlighted in magenta, was affected by strong line broadening in addition to a particularly large CSP, which is typical of fast-to-intermediate exchange kinetics between the ligand-free and ligand-bound states.

Despite the growing interest in autophagy inhibitors for cancer therapy and for autophagy-mediated degradation, few small molecules with potent and selective LC3/GABARAP binding have been reported. Here, we further investigated the structure-activity relationships of compound **1** against both LC3B and GABARAP. The best compounds have low micromolar IC_50_ values for inhibiting the interactions of these proteins with known peptide ligands. Because LC3/GABARAP proteins are highly conserved and expressed in every tissue at moderate to high levels,^45–47^ we anticipate that these compounds are likely not potent enough to make good candidates for autophagy inhibitors. Still, at micromolar potency they could be usable for autophagy-mediated targeted protein degradation. Importantly, any contributions from the documented ability of **1** to covalently bind E3 ligase DCAF11, which was not directly explored in this work, would have to be controlled for in any investigation of targeted protein degraders based on compound **1**. By contrast, the mechanism for the LC3B/GABARAP binding is unlikely to be covalent because these proteins lack cysteines. These observations suggest that the LC3/GABARAP binding may yet be separable from the Michael acceptor, either by saturation, substituting the Michael acceptor with a planar isostere, or by scaffold hopping to remove the Michael acceptor.

Overall, we provide definitive evidence that **1** binds GABARAP and inhibits the interactions of both LC3B and GABARAP with peptide ligands. The similarity of compound **1**-induced CSP patterns observed for GABARAPL2 (ref. 32) and GABARAP (this study) attests to the high conservation of the core LIR docking site and to the challenges of devising small-molecule ligands targeting individual family members. We also provide initial structure-activity relationships for selective recognition of LC3B and GABARAP by arylidene-indolinone compounds. Despite the ongoing controversy surrounding these compounds, our work suggests that compounds with similar scaffolds or similar pharmacophores could be viable LC3/GABARAP ligands for applications in cancer chemotherapy and targeted degradation of proteins, organelles, and protein aggregates. In such subsequent efforts, care should be taken to ensure solubility in all aqueous assay conditions and to address potential confounding effects due to DCAF11 binding. More broadly, this work adds to the growing evidence that these protein-protein interactions are highly likely to be druggable using orally bioavailable small molecules.

## ABBREVIATIONS

SPR,: surface plasmon resonance;
DMAP,: 4-dimethylaminopyridine;
EtOH,: ethanol;
NaBH_4_,: sodium borohydride;
DMF,: dimethylformamide;
LIR,: LC3-interacting region.

## ASSOCIATED CONTENT

### Supporting Information

The Supporting Information is available free of charge on the ACS Publications website.

Full experimental details, full compound characterization, complete binding assay data, and additional supporting data, PDF

## AUTHOR INFORMATION

### Author Contributions

A.L., H.B., and J.A.K. designed experiments. A.L., J.P., T.S., and B.T. prepared and characterized compounds. A.L. and H.B. measured binding affinities. P.N., A.Ü., O.H.W. and D.W. produced and interpreted NMR data and wrote the summary of the NMR results. A.L., H.B., and J.A.K. wrote the rest of the manuscript.

### Funding Sources

This work was funded by NIH GM148407 to J.A.K., and the Deutsche Forschungsgemeinschaft (DFG, German Research Foundation)–Project-ID 267205415–SFB 1208, project B02 (D.W.).

## ACKNOWLEDGMENT

The authors acknowledge access to the Jülich-Düsseldorf Biomolecular NMR Center.

## Notes

### Competing Interest Statement

The authors have declared no competing interest.

### Summary of Updates

data has been updated to account for low solubility of some compounds, which was not fully documented in previous version

