## Supplementary Materials for "Exploring Arylidene-Indolinone Ligands of Autophagy Proteins LC3B and GABARAP"

### Table Of Contents

1. **Experimental Section ........................................ S2**
    a. Protein Expression and Purification.......................... S2
    b. Calibration Curves and solubility data ...................... S2
    c. AlphaScreen Assay........................................ S4
    d. Beta-lactamase Assay........................................ S5
    e. E/Z Isomerization $^1$H NMR Experiment ..................... S6
    f. NMR Titration Experiments ...................................... S7
2. **General Information on Reaction Setup and Compound Characterization ........................................ S9**
3. **Synthesis Procedures and Characterization Data ........ S10**
4. **AlphaScreen Data ........................................ S16**
    a. Cross-Titration Experiments.................................... S16
    b. Assay Implementation ............................................ S17
5. **NMR Spectra for Novel Compounds.............................. S23**
6. **2D-NMR Data ........................................ S41**
7. **References ........................................ S42**

### 1. Experimental Section

**1a. *Protein expression and purification.*** Recombinant His-tagged GABARAP and LC3B were expressed in BL21 (DE3) *E. coli* transformed with pET15b expression plasmids encoding each protein fused with an N-terminal His tag. The LC3B expression plasmid was originally from Ma *et al.*,[1] and the GABARAP expression plasmid was designed to be identical but with the coding sequence for human GABARAP. Transformed cells were plated on ampicillin agar plates and incubated at 37 °C overnight. Individual colonies were picked and grown overnight shaking at 37 °C in 5 mL of LB culture medium with 1% ampicillin. Each 5 mL culture was then added to 1 L of LB culture medium with 1% ampicillin and incubated with shaking at 37 °C until the $OD_{600}$ measured ≥ 0.6. At this point, protein expression was induced by the addition of 1 mL of 1 mM Isopropyl β-D-1-thiogalactopyranoside (IPTG). Cells were then incubated for 3 hours at 37°C, then pelleted and stored at -80°C. To purify, cells were resuspended in a lysis buffer of 25 mM HEPES pH 7.3, 150 mM NaCl, 20 mM imidazole, 0.2% lysozyme, 1 protease inhibitor cocktail pellet (Roche), and 2.5 U/mL universal nuclease (Pierce). Resuspended cells were sonicated for 20 minutes in 10 second pulses, and lysed cells were spun down to separate the lysate and cellular debris. Clarified lysate was purified using batch affinity purification with HisPur Ni-NTA resin (ThermoFisher Scientific). Resin was incubated with the lysate at 4°C for one hour before washing with 25 mM HEPES pH 7.3, 150 mM NaCl, and 20 mM imidazole. Protein was then eluted from the resin in elution buffer (25 mM HEPES pH 7.3, 150 mM NaCl, and 500 mM imidazole). Protein purity and mass was evaluated by SDS-PAGE. Proteins were further purified by analytical size exclusion chromatology (Akta pure FPLC) if needed. Protein then underwent buffer exchange via a desalting column into a storage buffer (25 mM HEPES pH 7.3, 150 mM NaCl, 1 mM TCEP). Protein was then concentrated using Zeba desalting spin columns (Thermo Scientific), aliquoted, flash frozen in liquid $N_2$ and stored indefinitely at -80°C. Protein concentration was measured by absorbance at 280 nm (Thermo Scientific Nanodrop 1000).

**1b. Calibration Curves and Solubility Data.** Compounds were dissolved in DMSO to between 200 and 300 mM and serially diluted. The maximum absorbance wavelength

($\lambda_{max}$) of each compound was determined in DMSO using a Nanodrop microvolume spectrophotometer (Thermo Scientific) and concentration curves were created by measuring absorbance of serially diluted samples and plotting up to 6 concentrations. Extinction coefficients were derived from Beer's law using known concentrations and measured absorbance values. Once extinction coefficient in DMSO for a compound was determined, solubility in aqueous buffer could be determined routinely by removing an aliquot of saturated, aqueous compound, lyophilizing the water, resuspending in DMSO, and measuring absorbance. Maximum solubility in AlphaScreen buffer was determined by averaging the solubilities obtained in this manner from aqueous solutions prepared for each independent AlphaScreen assay trial.

**Table S1.** Maximum absorbance wavelengths, extinction coefficients, and maximum solubilities of prepared analogs. Each analog was tested in the AlphaScreen assay up to roughly 60% of its solubility limit. Errors represent standard errors of the mean from three independent trials.

| **Compound** | **$\lambda_{max}$ (nM)** | **Extinction Coefficient ($M^{-1}$ $cm^{-1}$)** | **Aqueous Solubility in AlphaScreen buffer (μM)** |
|---|---|---|---|
| **1** | 498 | 0.046 | 140 ± 6.3 |
| **2a** | 327 | 0.017 | 39.4 ± 2.3 |
| **2b** | 363 | 0.027 | 16.5 ± 1.2 |
| **2c** | 331 | 0.023 | 10.7 ± 2.0 |
| **2d** | 272 | 0.028 | 96.6 ± 10 |
| **2e** | 358 | 0.023 | 19.3 ± 4.2 |
| **2f** | 329 | 0.026 | 356 ± 18 |
| **2g** | 355 | 0.020 | 395 ± 16 |
| **2h** | 499 | 0.047 | 163 ± 14 |
| **2i** | 498 | 0.052 | 107 ± 10 |
| **2j** | 470 | 0.013 | 342 ± 29 |
| **2k** | 516 | 0.083 | 135 ± 28 |
| **2l** | 503 | 0.057 | 42.9 ± 7.3 |
| **2m** | 351 | 0.029 | 1.7 ± 0.1 |
| **2n** | 350 | 0.027 | 6.1 ± 3.5 |
| **2o** | 360 | 0.026 | 18.9 ± 4.6 |
| **2p** | 392 | 0.028 | 2.9 ± 0.8 |
| **2q** | 361 | 0.030 | 118 ± 33 |
| **2r** | 263 | 0.022 | 21.5 ± 3.6 |

**1c. *AlphaScreen Assay***

***Cross-Titration Experiments.*** A cross-titration experiment was performed on each bio-peptide/protein pair to establish ideal concentrations and assay conditions prior to testing of inhibitors in AlphaScreen, as described.[2] Each of GABARAP and LC3B were titrated from 300 nM to 10 nM, and each of bio-K1 and bio-FYCO1S were titrated from 30 nM to 1 nM. Conditions that minimized concentration while maintaining adequate signal-to-noise and a large difference in signal with varied concentration were chosen.

***Assay Implementation.*** Purified LC3B or GABARAP protein was diluted in assay buffer (25 mM HEPES pH 7.3, 150 mM NaCl, 0.1% Tween-20, 1 mg/mL BSA) to 200 nM and 100 nM (for LC3B) or 100 nm and 50 nM (for GABARAP). 2.5 µL of the more concentrated LC3B (200 nM) or GABARAP (100 nM) was added in duplicate to a white, 384-well polystyrene plate (AlphaPlate, Revvity) to wells intended for inhibitor compounds, and 5 µL dilute LC3B (100 nM) or GABARAP (50 nM) was added to control wells. Inhibitor compounds were prepared as stock solutions at 20 mM in DMSO. From these 20 mM stock solutions, 500 µM solutions of each compound in assay buffer with 7.5% DMSO were prepared in Eppendorfs. These solutions were then vortexed and centrifuged at 14800 rpm for one minute. The resulting supernatant was transferred to a new Eppendorf, re-vortexed and spun down again. This process, each time transferring only the supernatant, was repeated for each compound until no pellet formed following centrifugation, indicating that a saturated solution of the compound in assay buffer was achieved. These saturated solutions were then serially diluted in assay buffer with 7.5% DMSO. 15 µL of serially diluted compounds were added to wells with concentrated 2.5 µL protein. 5 µL of Acetyl-K1 and Acetyl-FYCO1S were added to wells with dilute 5 µL protein and thus were used as positive control inhibitors for GABARAP and LC3B, respectively. Plates were covered in foil, spun down at 1200 rpm for 3 minutes then incubated at room temperature for 45 minutes. Biotinylated K1 was diluted to 100 nM (concentrated) and 50 nM (dilute) in assay buffer, or biotinylated FYCO1S was diluted to 150 nM and 75 nM in assay buffer. 5 µL of the dilute bio-K1 or bio-FYCO1S was added to the control wells with GABARAP or LC3B, respectively, and 2.5 µL of the concentrated bio-K1 or Bio-FYCO1S was added to the wells with inhibitor compounds and GABARAP or LC3B, respectively. 2 blank wells containing 10 µL assay buffer and 5 µL assay buffer + 7.5% DMSO, as well as 2 no-inhibitor control wells containing 5 µL dilute protein, 5 µL dilute bio-peptide, and 5 µL assay buffer + 7.5% DMSO were included in each plate. Again, plates were covered in foil, spun down, and incubated for 45 minutes. AlphaScreen streptavidin donor beads and nickel chelate acceptor beads (Revvity) were each diluted to 200 µg/mL (concentrated) and 100 µg/mL (dilute) in assay buffer. 5 µL dilute acceptor beads were added to each control well, and 2.5 µL concentrated acceptor beads were added to each inhibitor compound well. Immediately following this and in the dark, 5 µL dilute donor beads were added to each control, and 2.5 µL concentrated donor beads were added to each inhibitor compound well. Plates were covered, spun down again, and incubated for 1 hour before being read on a plate reader (Tecan Spark, AlphaScreen method, excitation at 680 nm and emission at 520-620 nm). Using GraphPad Prism 10 software, data were normalized using blank wells as 0% bound and no inhibitor control wells as 100% bound, then fit with $IC_{50}$ curves. Average $IC_{50}$ and standard errors of the mean were calculated based on at least three independent trials.

**1d*. Beta-lactamase Assay.*** The Beta-lactamase inhibitor screening kit was purchased from Sigma-Aldrich. Reagents were warmed to room temperature and briefly centrifuged before opening. Beta-lactamase was reconstituted in 220 µL of the beta-lactamase buffer and aliquoted into several microcentrifuge tubes. Aliquots not used immediately were stored at -20 °C. In a new microcentrifuge tube, 3 µL of inhibitor control stock solution was added to 47 µL of the beta-lactamase buffer. To a third microcentrifuge tube, 9 µL of nitrocefin was added to 261 µL of assay buffer (25 mM HEPES pH 7.3, 150 mM NaCl, 0.1% Tween-20, 1 mg/mL BSA). Serial dilutions of compound **1** was prepared in a 96-well plate using the beta-lactamase buffer provided. Final concentrations of compounds in well 1 were 25 µM and were diluted by 1:3 to give 5 dilutions.

To a clear, polystyrene, flat-bottom 96 well plate, 100 µL of assay buffer was added to wells A1-2 for a blank (Table S1). For Reference 1 (100% enzyme activity control), to wells A4-5 was added 20 µL of assay buffer, 50 µL of enzyme solution, and 30 µL of

nitrocefin solution. For Reference 2 (positive control inhibitor), to wells A7-8 was added 20 μL of inhibitor control solution, 50 μL of enzyme solutions, and 30 μL of nitrocefin. To wells 1-5 for B-G was added 20 μL of the corresponding compound serial dilutions. For the blank serial dilution wells (B1-5, D1-5, and G1-5) was added 80 μL of assay buffer. To the serial dilution wells (C1-5, E1-5, and F1-5) was added 50 μL of the enzyme solution. The plate was then allowed to incubate for 15 minutes at room temperature before adding 30 μL of nitrocefin solutions to the serial dilution wells (C1-5, E1-5, and F1-5). Upon addition, the solution was pipetted up and down three times to mix.
The absorbance of each well was measured at 490 nm on a plate reader (Tecan Spark) after 15, 30, and 60 minutes of incubation at room temperature. The values from the blank wells were subtracted from the other values before plotting (Fig. S1).

**Figure S1.** Beta-lactamase Assay Inhibition Data for Compound **1**

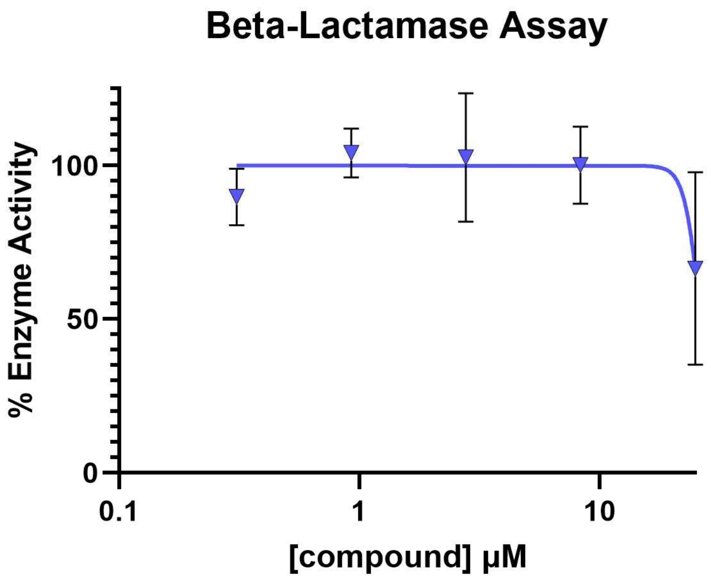


**1e.** ***E/Z Isomerization $^1$H NMR Experiment.*** To determine the E/Z ratios of the compounds and whether they isomerize in DMSO at room temperature, $^1$H NMR was taken at time 0, 2 hours, and 6 hours (Fig. S4, compound **2c**). As shown in the Fig. S4, the E isomer is completely racemized by hour 6 in DMSO. Though we are only showing one example, all compounds exhibited similar racemization patterns.

**Figure S2.** $^{1}H$ NMR Isomerization Time Trial of Compound **2c** in DMSO

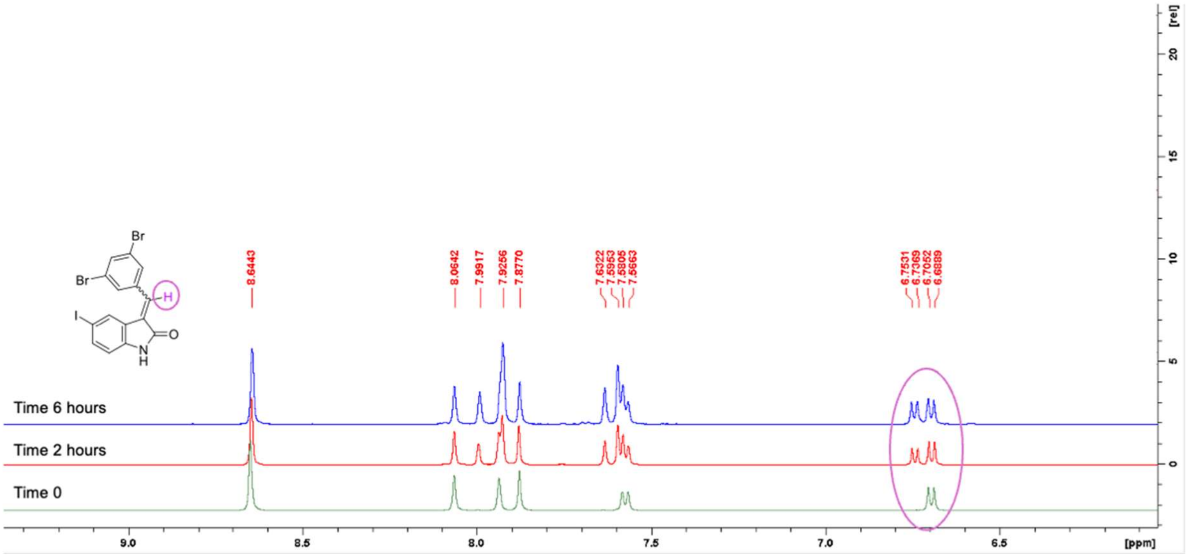


### 1f. *NMR Titration Experiments*

**Protein expression and purification for NMR spectroscopy.** [U-$^{15}N$] GST-GABARAP protein was purified from *E. coli* BL21(DE3) transformed with pGEX4T2-GABARAP and cultivated in M9 minimal medium containing $^{15}NH_4Cl$ (1 g/L). Gene expression was induced with 1 mM IPTG at an $OD_{600nm}$ of 0.7 and allowed to proceed for 20 hours at 25 °C; afterwards cells were harvested by centrifugation at 3000 × g for 30 min at 4 °C. The GST fusion protein was purified from the soluble extract by affinity chromatography using Glutathione Sepharose 4B (GE Healthcare). Cleavage with thrombin yielded a 119-aa protein carrying an N-terminal glycine-serine extension in addition to the native residues of GABARAP. Afterwards, the sample was applied to a Hiload 26/60 Superdex 75 preparatory grade size exclusion column equilibrated with 25 mM $NaH_2PO_4$/$Na_2HPO_4$, 100 mM NaCl, 100 mM KCl, 50 µM EDTA, pH 6.9. The protein was eluted in the same buffer, snap frozen in liquid $N_2$ and stored at -80°C until further use.

**NMR titration experiments.** In order to enhance the solubility of the ligand, the titration was performed in a mixture of water and DMSO. To this end, a sample of 200 µl 200 µM [U-$^{15}N$] GABARAP with 20 mM sodium phosphate, 80 mM NaCl, 80 mM KCl, 40 µM EDTA in 10% (v/v) DMSO, 10% (v/v) $D_2O$, 80% (v/v) $H_2O$, pH 6.9 in a 3 mm NMR tube was titrated with a stock solution of 5.0 mM GW5074 with 13 mM sodium phosphate, 50 mM NaCl, 50 mM KCl, 25 µM EDTA in 50% (v/v) DMSO, 50% (v/v) $H_2O$, pH 6.9, resulting in final DMSO concentrations below 14% (v/v) up to a ligand:protein stoichiometry of 5:2. To verify the integrity of the tertiary structure under these conditions and to identify the resonances affected by changes in DMSO concentration, we performed a control titration of a sample of 200 µl 200 µM [U-$^{15}N$] GABARAP with 23 mM sodium phosphate, 90 mM NaCl, 90 mM KCl, 45 µM EDTA in 10% (v/v) $D_2O$, 90% (v/v) $H_2O$, pH 6.9 with increasing amounts of DMSO up to a final concentration of 14% (v/v).

Titrations were monitored by recording 2D [$^{1}H$,$^{15}N$] HSQC spectra[3] at a temperature of 25.0°C on a Bruker AVANCE III HD 700 MHz NMR spectrometer equipped with a cryogenically cooled triple resonance probe with z axis pulsed field gradient capabilities. The sample temperature was calibrated using methanol-$d_4$.[4] The $H_2O$ and DMSO resonances were suppressed by gradient coherence selection, with quadrature detection in the indirect $^{15}N$ dimension achieved by the echo-antiecho method[5,6]. A WALTZ-16 sequence with a field strength of 1.1 kHz was employed for $^{15}N$ decoupling during acquisition.[7] 1024 (192) complex data points were acquired with a spectral width of 16 ppm (29.0 ppm) in the $^{1}H$ ($^{15}N$) dimension. NMR spectra were processed using NMRPipe and NMRDraw[8] and analyzed with NMRViewJ.[9] $^{1}H$ chemical shifts were referenced with respect to external DSS (2,2-dimethylsilapentane-5-sulfonic acid) in $D_2O$ and $^{15}N$ chemical shifts were referenced indirectly.[10] The control titration was used to propagate the sequence-specific resonance assignments of GABARAP in aqueous buffer[11] to the water/DMSO mixtures used in this study. For chemical shift perturbation (CSP) analysis, $^{1}H$ chemical shift changes, $\Delta\delta(^{1}H)$, and $^{15}N$ chemical shift changes, $\Delta\delta(^{15}N)$, in units of ppm were combined according to the following weights:

$$\Delta\delta(^1\mathrm{H}, ^{15}\mathrm{N}) = \sqrt{\left(\Delta\delta(^1\mathrm{H})\right)^2 + \left(0.2 \times \Delta\delta(^{15}\mathrm{N})\right)^2}$$

Chemical shift perturbations exceeding twice the root-mean-square CSP (calculated over all assigned resonances not affected by large chemical shift changes) were considered significant. With the exception of the side-chain amide resonances of N81, which were excluded from further analysis, none of the amide resonances exhibiting a significant CSP upon titration with GW5074 was among the resonances significantly affected by the DMSO control titration (K6, I41, D45, H69, N81 side-chain, V83, A89, Q96, E100, Y115, G116).

### 2. General Information on Reaction Setup and Compound Characterization

***Reaction setup.*** Reactions were conducted in oven-dried glassware equipped with tightly fitted rubber septa and under a positive pressure of dry nitrogen. Reagents and solvents were handled by using standard syringe techniques. Unless stated otherwise, all yields refer to isolated products after flash column chromatography.

***NMR Spectroscopy.*** $^1$H NMR spectra of compounds were acquired using a Bruker Biospin 500 MHz Avance III NMR spectrometer and calibrated using the solvent signal (DMSO-$d_6$ 2.50 ppm). Multiplicities were determined using TopSpin software. $^{13}$C NMR spectra of compounds were acquired using a Bruker Biospin 500 MHz Avance III NMR spectrometer and calibrated using the solvent signal (DMSO-$d_6$ 39.51 ppm). Chemical shifts ($\delta$) are reported in parts per million (ppm) and coupling constants (*J*) are measured in hertz (Hz) and calculated using Topspin software. The following abbreviations were used to describe multiplicities: s = singlet, d = doublet, t = triplet, m = multiplet.

***Mass Spectroscopy.*** Electrospray ionization mass spectra were acquired using a Thermo Finnigan LTQ mass spectrometer.

***Solvents/chemicals.*** All solvents and reagents were used as received from the manufacturer.

### 3. Synthesis Procedures and Characterization Data

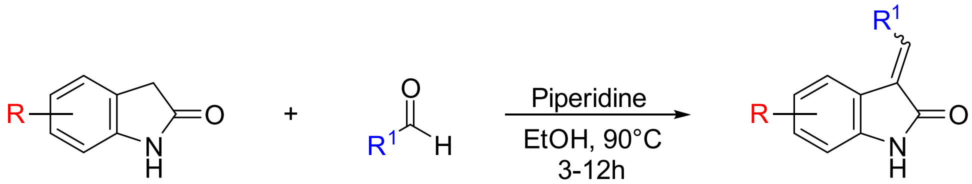


Compounds were synthesized and purified using previously reported procedures[12]. Compounds were isolated as a mixture of E/Z isomers or isolated as one isomer that isomerized to a roughly 50:50 mixture once dissolved in DMSO at room temperature (Fig. S2).

3-benzylidene-5-iodoindolin-2-one **2a**

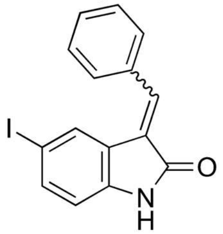


Yellow solid; yield: 54%. **$^1$H NMR** (500 MHz, DMSO-$d_6$): δ 10.83 (broad, 1H), 8.39 (d, *J* = 4 Hz, 1H), 8.10 (s, 1H), 7.92 (s, 1H), 7.69 (m, 4H), 7.54 (m, 4H), 6.74 (d, *J* = 8.15 Hz, 1H), 6.67 (d, *J* = 8 Hz, 1H) ppm; **$^{13}$C NMR (**125 MHz, DMSO-$d_6$**)** δ 167.9, 166.5, 142.5, 140.2, 138.5, 138.2, 137.3, 136.9, 134.1, 133.8, 132.1, 130.7, 130.1, 130.0, 129.1, 128.7, 128.2, 128.1, 127.5, 126.7, 125.4, 123.5, 112.5, 111.7, 83.9, 83.6 ppm; **HRMS** (ESI): calcd for $C_{15}H_{11}INO^+$ [M+H$^+$]: 347.988; found: 347.981

3-(4-hydroxybenzylidene)-5-iodoindolin-2-one **2b**

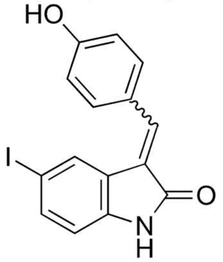


Yellow solid; yield: 56%. **$^1$H NMR** (500 MHz, DMSO-$d_6$) δ 10.67 (s, 1H), 10.31 (broad, 1H), 8.43, (d, *J* = 7.5 Hz, 2H), 8.04 (s, 1H), 7.82 (s, 1H), 7.48 (d, *J* = 7.24 Hz, 1H), 6.87 (d, *J* = 7.59 Hz, 2H), 6.66 (d, *J* = 7.63 Hz, 1H) ppm; **$^{13}$C NMR (**125 MHz, DMSO-$d_6$**)** δ 190.9, 168.3, 166.8, 160.5, 159.6, 142.0, 139.4, 139.2, 138.2, 137.4, 135.8, 135.2, 132.1, 131.9, 129.7, 128.3, 127.3, 125.5, 124.6, 124.0, 123.6, 121.4, 115.8, 115.6, 115.3, 112.3, 111.4, 83.8, 83.6 ppm; **HRMS** (ESI): calcd for $C_{15}H_{11}INO_2^+$ [M+H$^+$]: 363.983; found: 363.988.

3-(3,5-dibromobenzylidene)-5-iodoindolin-2-one **2c**

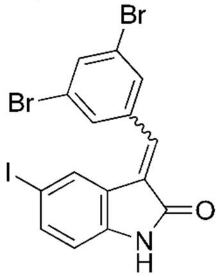

Yellow solid; yield: 79%. **$^1$H NMR** (500 MHz, DMSO-$d_6$) δ 10.83 (s, 1H), 8.61 (s, 2H), 8.03 (s, 1H), 7.87 (d, *J* = 29.5 Hz, 2H), 7.54 (d, *J* = 8.05 Hz, 1H) , 6.66 (d, *J* = 8.10 Hz, 1H) ppm; **$^{13}$C NMR (**125 MHz, DMSO-$d_6$**)** δ 166.3, 140.7, 137.8, 137.5, 134.7, 134.6, 133.2, 128.6, 128.0, 126.8, 122.1, 112.0, 84.2 ppm; **HRMS** (ESI): calcd for $C_{15}H_9Br_2INO^+$ [M+H$^+$]: 503.809; found: 503.800

3-(3,5-dibromo-2-hydroxybenzylidene)-5-iodoindolin-2-one **2d**

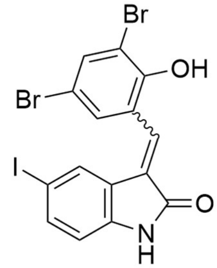

Yellow solid; yield: 65%. **$^1$H NMR** (500 MHz, DMSO-$d_6$) δ 10.79 (s, 1H), 10.43 (s, 1H), 8.44 (s, 1H), 8.02 (s, 1H), 7.85 (d, *J* = 22.78, 2H), 7.57 (s, 1H), 6.71 (s, 1H) ppm; **$^{13}$C NMR (**125 MHz, DMSO-$d_6$**)** δ 167.5, 166.4, 151.9, 142.5, 140.5, 138.3, 137.5, 136.1, 135.8, 133.2, 131.4, 131.3, 131.1, 130.9, 128.6, 128.1, 126.8, 125.6, 124.8, 123.3, 113.3, 112.7, 112.5, 111.9, 110.7, 84.1, 83.8 ppm; **HRMS** (ESI): calcd for $C_{15}H_9Br_2INO_2^+$ [M+H$^+$]: 519.804; found: 519.806.

5-iodo-3-(4-methoxybenzylidene)indolin-2-one **2e**

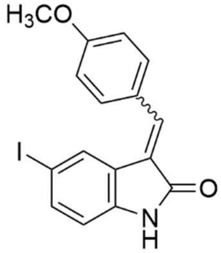

Yellow solid; yield: 51%. **$^1$H NMR** (500 MHz, DMSO-$d_6$) δ 10.69 (s, 1H), 7.88 (s, 1H), 7.70 (d, *J* = 8.20 Hz, 2H), 7.64 (s, 1H), 7.55 (d, *J* = 8.10 Hz, 1H), 7.12 (d, *J* = 8.10 Hz, 2H), 6.73 (d, *J* = 8.10 Hz, 1H), 3.87 (s, 3H) ppm; **$^{13}$C NMR (**125 MHz, DMSO-$d_6$**)** δ 168.7, 161.4, 142.6, 138.2, 138.1, 132.1, 130.3, 126.7, 125.0, 124.3, 114.8, 112.9, 84.2, 56.0 ppm; **HRMS** (ESI): calcd for $C_{16}H_{13}INO_2^+$ [M+H$^+$]:377.999; found: 377.990

4-((5-iodo-2-oxoindolin-3-ylidene)methyl)benzoic acid **2f**

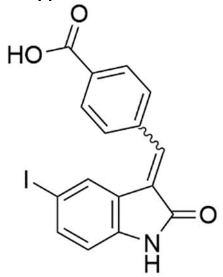

Yellow solid; **yield**: 62%. **$^{1}$H NMR** (500 MHz, DMSO-d$_6$) δ 13.20 (s, 1H), 10.78 (s, 1H), 8.06 (d, *J* = 6.40 Hz, 2H), 7.78 (d, *J* = 6.60 Hz, 2H), 7.69 (s, 1H), 7.63 (s, 1H), 7.56 (d, *J* = 7.00 Hz, 1H), 6.72 (d, *J* = 7.50 Hz, 1H) ppm; **$^{13}$C NMR (**125 MHz, DMSO-d$_6$**)** δ 168.2, 167.2, 143.2, 139.1, 139.0, 136.5, 132.1, 130.9, 130.1, 129.9, 128.4, 123.7, 113.1, 84.4 ppm; **HRMS** (ESI): calcd for $C_{16}H_{10}INO_3^+$ [M+H$^+$]: 391.164.

3-(3,5-dibromo-4-hydroxybenzylidene)indolin-2-one **2g**

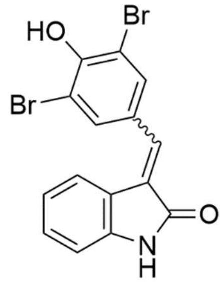

Yellow solid; yield: 52%. **$^{1}$H NMR** (500 MHz, DMSO-d$_6$): δ 10.34 (s, 1H), 8.73 (s, 2H), 8.19 (d, *J* = 6.80 Hz, 2H), 7.53 (d, *J* = 7.50 Hz, 1H), 7.46 (s, 1H), 7.05 (t, *J* = 7.57 Hz, 1H), 6.90 (m, *J* = 6.31 Hz, 3H), 6.77 (d, *J* = 7.60 Hz, 1H) ppm; **$^{13}$C NMR (**125 MHz, DMSO-d$_6$**)** δ 189.3, 176.3, 168.4, 167.2, 143.6, 140.6, 136.0, 133.9, 133.6, 133.1, 130.3, 128.9, 128.6, 127.4, 126.0, 125.7, 124.8, 124.3, 121.1, 121.1, 119.6, 112.1, 111.7, 111.0, 109.4, 109.0 ppm; **HRMS** (ESI): calcd for $C_{15}H_{10}Br_2NO_2^+$ [M+H$^+$]: 393.907; found: 393.912.

3-(3,5-dibromo-4-hydroxybenzylidene)-6-iodoindolin-2-one **2h**

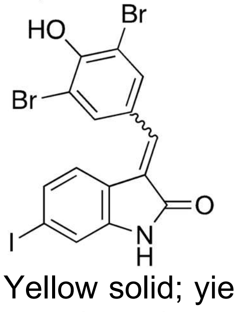

Yellow solid; yield: 47%. **$^{1}$H NMR** (500 MHz, DMSO-d$_6$): δ 10.70 (s, 1H), 10.62 (s, 1H), 7.91 (s, 2H), 7.55 (s, 1H), 7.27 (dd, $J_1$ = 8.00, $J_2$ = 7.95, 2H), 7.21 (s, 1H) ppm; **$^{13}$C NMR (**125 MHz, DMSO-d$_6$**)**: δ 168.6, 167.4, 153.3, 152.6, 144.7, 142.3, 136.7, 135.5, 134.6, 133.7, 130.3, 130.2, 129.0, 127.2, 125.6, 125.1, 123.9, 121.9, 120.9, 119.1, 118.2, 112.3, 111.6, 96.4, 94.6 ppm; **HRMS** (ESI): calcd for $C_{15}H_9Br_2INO_2^+$ [M+H$^+$]: 519.804; found: 519.811.

5-bromo-3-(3,5-dibromo-4-hydroxybenzylidene)indolin-2-one **2i**

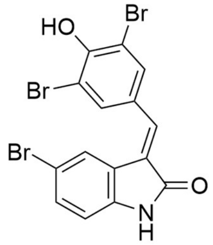

Yellow solid; yield: 32%. **$^{1}$H NMR** (500 MHz, DMSO-$d_6$): δ 10.81 (s, 1H), 10.77 (s, 1H), 8.80 (s, 2H), 7.90 (s, 1H), 7.84 (s, 1H), 7.37 (d, *J* = 8.13, 1H), 6.80 (d, *J* = 8.26, 1H) ppm; **$^{13}$C NMR (**125 MHz, DMSO-$d_6$**)**: δ 168.5, 167.4, 153.6, 152.8, 142.6, 140.1, 136.9, 136.4, 135.5, 133.9, 132.9, 131.4, 128.7, 128.6, 127.8, 127.0, 125.1, 125.0, 123.4, 122.8, 113.6, 113.1, 112.6, 112.3, 112.3, 111.8, 111.6 ppm; **HRMS** (ESI): calcd for $C_{15}H_8Br_3NO_2^+$ [M+H$^+$]: 471.818; found: 471.872

3-(3,5-dibromo-4-hydroxybenzylidene)-6-methylindolin-2-one **2j**

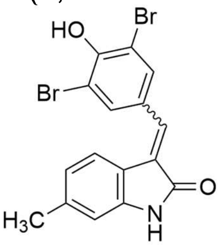

Yellow solid; yield: 41%. **$^{1}$H NMR** (500 MHz, DMSO-$d_6$): δ 10.62 (s, 1H), 7.93 (s, 1H), 7.87 (s, 2H), 7.44 (s, 1H), 7.21 (d, *J* = 7.70, 1H), 6.70 (s, 1H), 6.69 (d, *J* = 8.15, 1H) ppm; **$^{13}$C NMR (**125 MHz, DMSO-$d_6$**)**: δ 168.9, 144.1, 141.6, 139.4, 134.3, 131.3, 131.1, 130.2, 123.2, 122.6, 122.4, 118.2, 111.5, 22.0 ppm; **HRMS** (ESI): calcd for $C_{16}H_{11}Br_2NO_2^+$ [M+H$^+$]: 407.923; found: 408.079

Methyl 3-(3,5-dibromo-4-hydroxybenzylidene)-2-oxoindoline-6-carboxylate **2k**

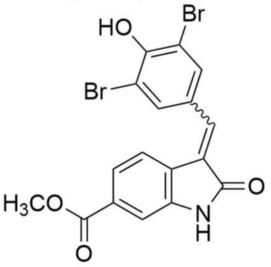

Yellow solid; yield: 65%. **$^{1}$H NMR** (500 MHz, DMSO-$d_6$): δ 10.89 (s, 1H), 10.83 (s, 1H), 8.85 (s, 2H), 7.89 (s, 1H), 7.78 (d, *J* = 7.85, 1H), 7.64 (d, *J* = 6.75, 1H) 7.35 (s, 1H), 3.86 (s, 3H) ppm; **$^{13}$C NMR (**125 MHz, DMSO-$d_6$**)**: δ 168.7, 167.5, 166.5, 166.2, 153.7, 152.9, 143.6, 141.0, 137.6, 137.1, 136.7, 134.0, 130.9, 130.2, 129.8, 128.8, 128.7, 127.1, 125.7, 125.4, 123.0, 122.4, 120.1, 112.3, 111.6, 110.6, 110.0, 52.8, 52.7 ppm; **HRMS** (ESI): calcd for $C_{17}H_{11}Br_2NO_4^+$ [M+H$^+$]: 451.913; found: 451.932

5,6-dichloro-3-(3,5-dibromo-4-hydroxybenzylidene)indolin-2-one **2l**

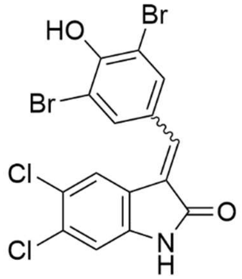


Yellow solid; **yield**: 32%. **$^1$H NMR** (500 MHz, DMSO-d$_6$) δ 10.90 (s, 1H), 10.75 (s, 1H), 7.96 (s, 2H), 7.61 (d, *J* = 4.65 Hz, 2H), 7.07 (s, 1H) ppm; **$^{13}$C NMR (**125 MHz, DMSO-d$_6$**)** δ 168.6, 167.4, 153.8, 140.6, 137.2, 137.0, 133.9, 130.8, 128.6, 126.3, 124.3, 123.8, 121.9, 121.8, 112.4, 112.1, 111.7, 111.4 ppm; **HRMS** (ESI): calcd for $C_{15}H_7Br_2Cl_2NO_2$ [M$^+$H$^+$]: 461.821; found: 461.904.

5-iodo-3-((4'-(trifluoromethyl)-[1,1'-biphenyl]-4-yl)methylene)indolin-2-one **2m**

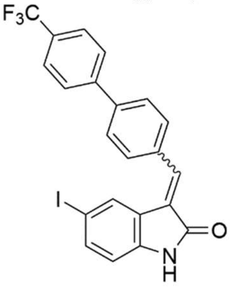


Yellow solid; yield: 44%. **$^1$H NMR** (500 MHz, DMSO-d$_6$) δ 10.78 (s, 1H), 8.54 (d, *J* = 8.10 Hz, 1H), 8.00 (d, J = 7.90 Hz, 2H), 7.95 (d, J = 8.05 Hz, 2H), 7.85 (m, J = 7.12 Hz, 5H), 7.72 (s, 1H), 7.57 (d, J = 7.55 Hz, 1H), 7.54 (d, J = 8.45 Hz, 1H), 6.75 (d, J = 8.10 Hz, 1H), 6.68 (d, J = 8.05 Hz, 1H) ppm; **$^{13}$C NMR (**125 MHz, DMSO-d$_6$**)** δ 168.4, 143.5, 143.1, 140.8, 140.4, 138.9, 137.1, 134.7, 133.4, 130.8, 130.7, 128.7, 128.1, 128.1, 127.8, 127.3, 126.4, 126.4, 125.9, 123.9, 123.7, 113.1, 112.2, 84.3 ppm; **HRMS** (ESI): calcd for $C_{22}H_{14}F_3INO^+$ [M+H$^+$]: 492.007; found: 492.001.

5-iodo-3-(thiazol-5-ylmethylene)indolin-2-one **2n**

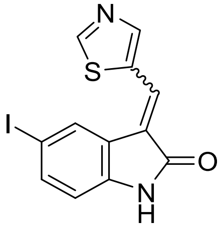


Yellow solid; yield: 72%. **$^1$H NMR** (500 MHz, DMSO-d$_6$) δ 10.83 (s, 1H), 9.27 (s, 1H), 8.46 (s, 1H), 8.36 (s, 1H), 8.07 (s, 1H), 7.53 (d, *J* = 8.05 Hz, 1H), 6.70 (d, *J* = 8.15 Hz, 1H) ppm; **$^{13}$C NMR (**125 MHz, DMSO-d$_6$**)** δ 167.2, 161.3, 153.4, 140.8, 137.6, 132.3, 128.7, 127.0, 122.8, 112.6, 84.7 ppm; **HRMS** (ESI): calcd for $C_{12}H_8IN_2OS^+$ [M+H$^+$]: 354.940; found: 354.942.

5-iodo-3-((4-methylthiazol-5-yl)methylene)indolin-2-one **2o**

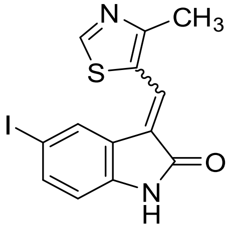


Yellow solid; **yield**: 57%. **$^1$H NMR** (500 MHz, DMSO-d$_6$) δ 10.74 (s, 1H), 9.13 (s, 1H), 8.30 (s, 1H), 8.04 (s, 1H), 7.50 (d, *J* = 7.99 Hz, 1H), 6.67 (d, *J* = 8.10 Hz, 1H), 2.72 (s, 3H) ppm; **$^{13}$C NMR (**125 MHz, DMSO-d$_6$**)** δ 167.3, 161.9, 158.6, 140.5, 137.3, 128.9, 127.2, 126.6, 125.6, 121.9, 112.3, 84.6, 16.7 ppm; **HRMS** (ESI): calcd for $C_{13}H_{10}IN_2OS^+$ [M+H$^+$]: 368.955; found: 368.946.

3-((1H-pyrrol-2-yl)methylene)-5-iodoindolin-2-one **2p**

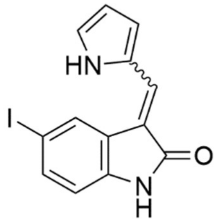


Yellow solid; yield: 24%. **$^1$H NMR** (500 MHz, DMSO-d$_6$) δ 13.28 (s, 1H), 11.00 (s, 1H), 8.02 (s, 1H), 7,89 (s, 1H), 7.45 (d, *J* = 8.05 Hz, 1H), 7.40 (s, 1H), 6.85 (s, 1H), 6.73 (d, *J* = 8.06 Hz, 1H), 6.39 (s, 1H) ppm; **$^{13}$C NMR (**125 MHz, DMSO-d$_6$**)** δ 168.4, 138.0, 135.6, 134.4, 132.4, 129.3, 127.7, 127.6, 126.5, 126.2, 120.9, 114.9. 111.6, 111.5, 111.2, 84.1 ppm; **HRMS** (ESI): calcd for $C_{13}H_{10}IN_2O^+$ [M+H$^+$]: 336.983; found: 336.989.

3-((1H-imidazol-5-yl)methylene)-5-iodoindolin-2-one **2q**

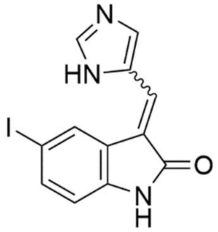


Yellow solid; yield: 26%. **$^1$H NMR** (500 MHz, DMSO-d$_6$) δ 13.60 (s, 1H), 11.08 (s, 1H), 8.03 (s, 2H), 7.97 (s, 1H), 7.61 (s, 1H), 7.48 (d, *J* = 7.45 Hz, 1H), 6.72 (d, *J* = 7.50 Hz, 1H) ppm; **$^{13}$C NMR (**125 MHz, DMSO-d$_6$**)** δ 168.9, 140.5, 139.7, 139.6, 136.4, 128.6, 128.1, 127.6, 124.9, 119.0, 112.6, 85.1 ppm; **HRMS** (ESI): calcd for $C_{12}H_9IN_3O^+$ [M+H$^+$]: 337.978; found: 337.970.

3-(imidazo[1,2-a]pyridin-5-ylmethylene)-5-iodoindolin-2-one **2r**

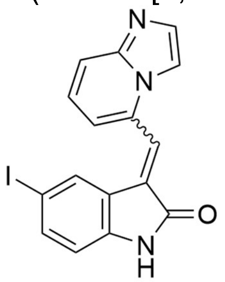

Orange solid; **yield**: 38%. **$^1$H NMR** (500 MHz, DMSO-$d_6$) δ 10.92 (s, 1H), 8.42 (m, 1H), 7.93 (s, 1H), 7.80 (d, *J* = 8.75 Hz, 1H), 7.74 (m, 2H), 7.60 (d, *J* = 7.60 Hz, 1H), 7.43 (s, 2H), 6.77 (d, *J* = 8.10 Hz, 1H) ppm; **$^{13}$C NMR (**125 MHz, DMSO-$d_6$**)** δ 168.0, 166.5, 145.4, 145.2, 143.6, 141.4, 139.7, 138.8, 134.7, 134.1, 132.0, 131.9, 131.1, 130.2, 129.3, 126.9, 126.2, 125.8, 124.0, 123.9, 123.2, 119.5, 119.1, 117.4, 115.1, 113.2, 112.8, 112.6, 112.4, 84.8, 84.48 ppm; **HRMS** (ESI): calcd for $C_{16}H_{10}IN_3O^+$ [M+H$^+$]: 387.994; found: 388.006.

### 4. AlphaScreen Data

#### 4a. Cross-Titration Experiments

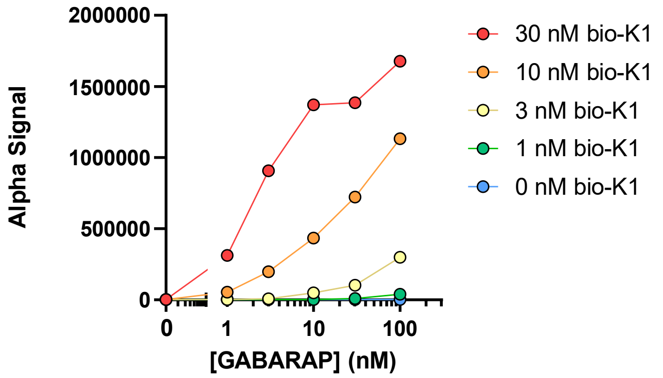


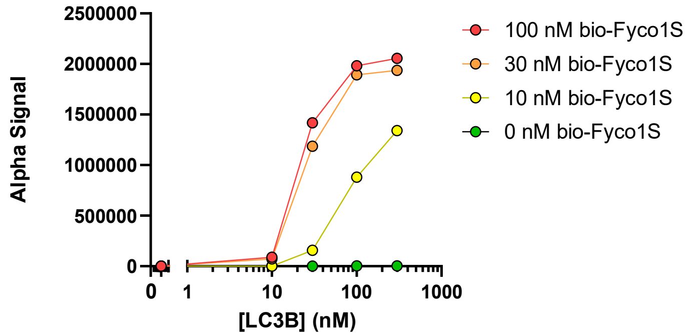


Representative cross-titration experiments for (a) bio-K1 and GABARAP and (b) bio-Fyco1S and LC3B.

### 4b. Assay Implementation

This section shows all relevant AlphaScreen data and curve fits for small molecules that displayed inhibiting interactions of recombinant GABARAP and LC3B to their respective peptide ligands. Two independent trials are shown for molecules that show no binding or binding only above 100 μM. Three independent trials are shown for compounds that show binding under 100 μM. Acetylated peptide FYCO-1S and K1 were used as positive controls for LC3B and GABARAP, respectively.

GABARAP LC3B

**FYCO-1S**

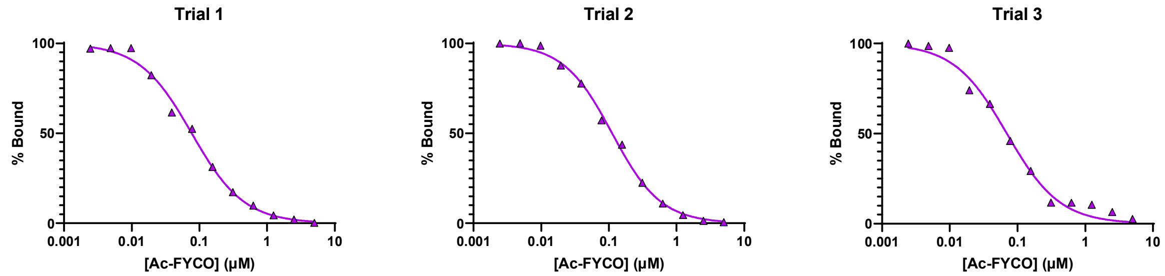


**K1**

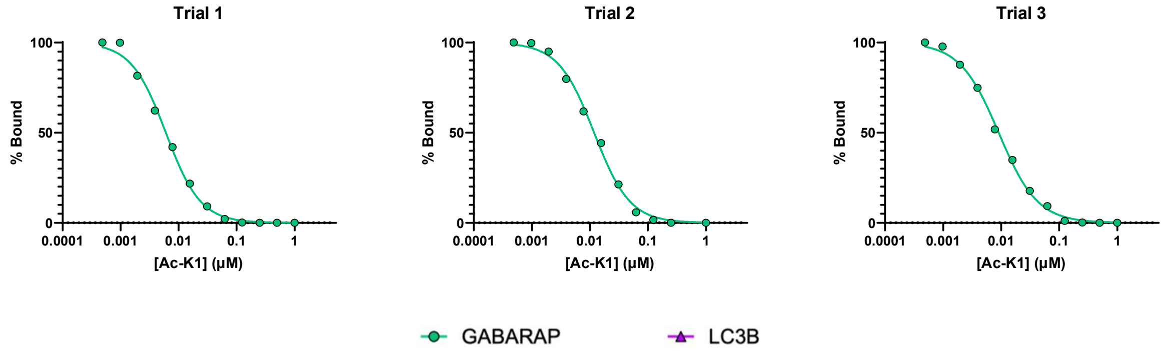

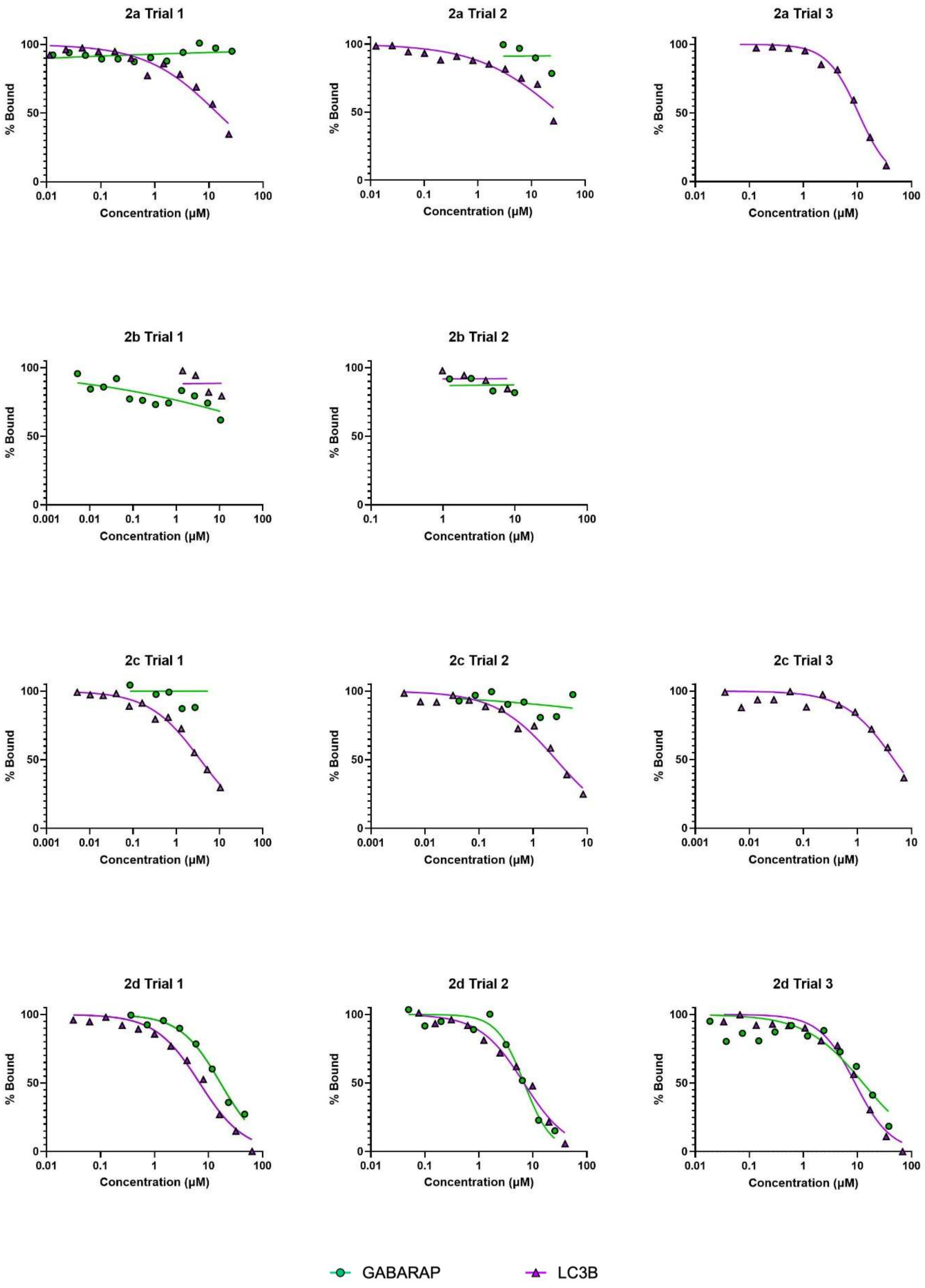
2a Trial 1
2a Trial 2
2a Trial 3
2b Trial 1
2b Trial 2
2c Trial 1
2c Trial 2
2c Trial 3
2d Trial 1
2d Trial 2
2d Trial 3
% Bound
Concentration (µM)
GABARAP
LC3B

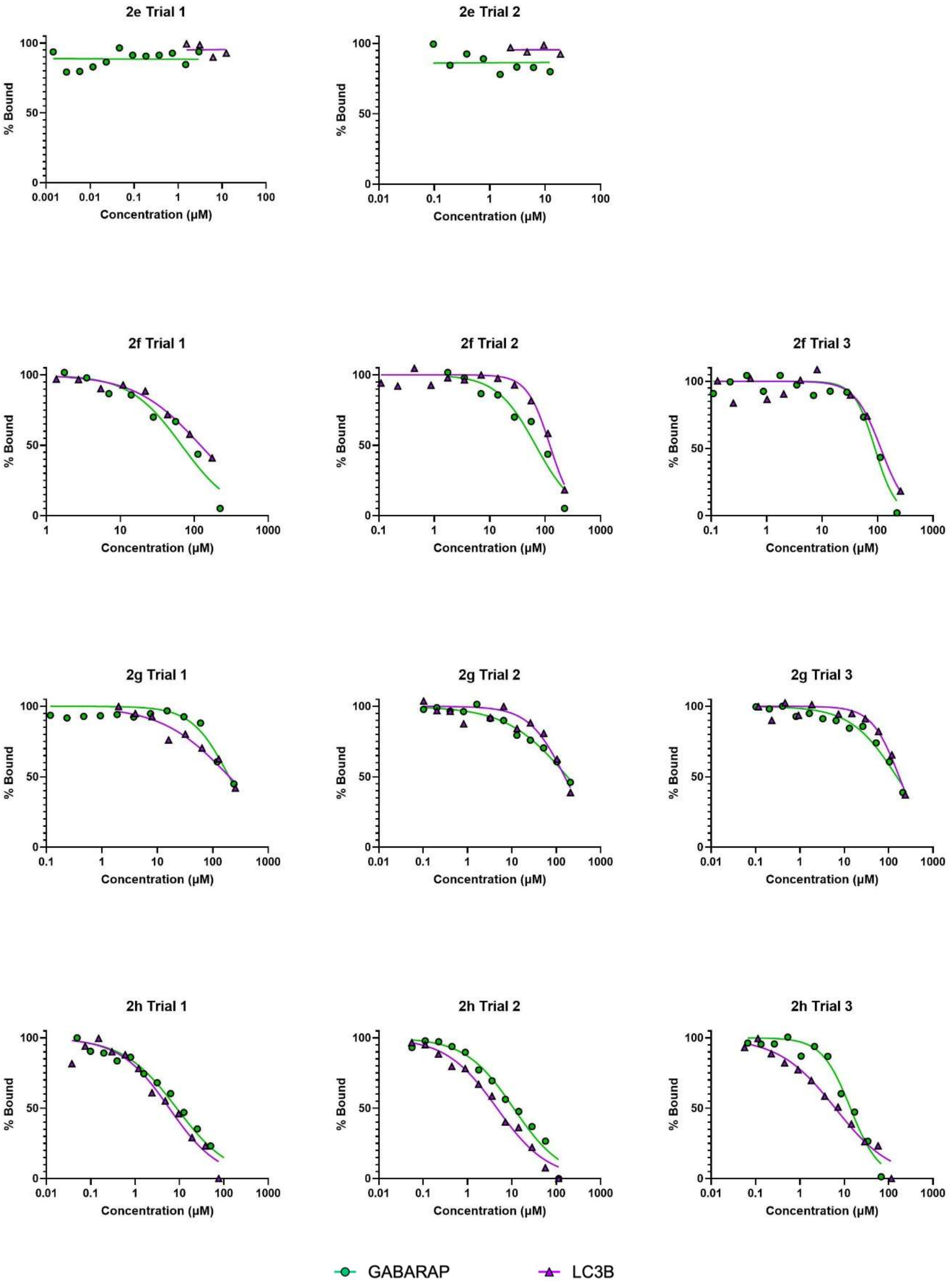
2e Trial 1
2e Trial 2
2f Trial 1
2f Trial 2
2f Trial 3
2g Trial 1
2g Trial 2
2g Trial 3
2h Trial 1
2h Trial 2
2h Trial 3
% Bound
Concentration (μM)
GABARAP
LC3B

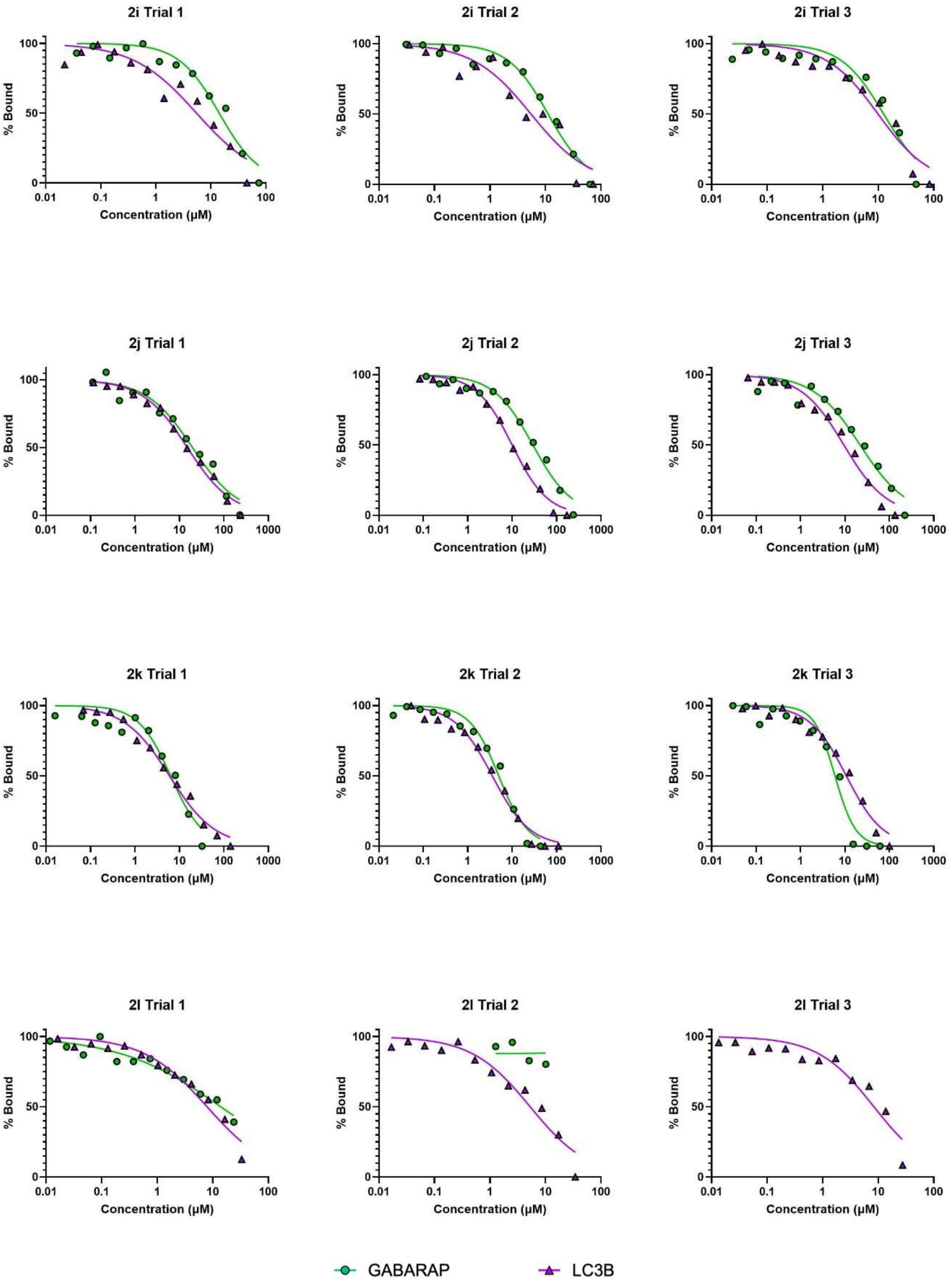
2i Trial 1
2i Trial 2
2i Trial 3
2j Trial 1
2j Trial 2
2j Trial 3
2k Trial 1
2k Trial 2
2k Trial 3
2l Trial 1
2l Trial 2
2l Trial 3
% Bound
Concentration (μM)
GABARAP
LC3B

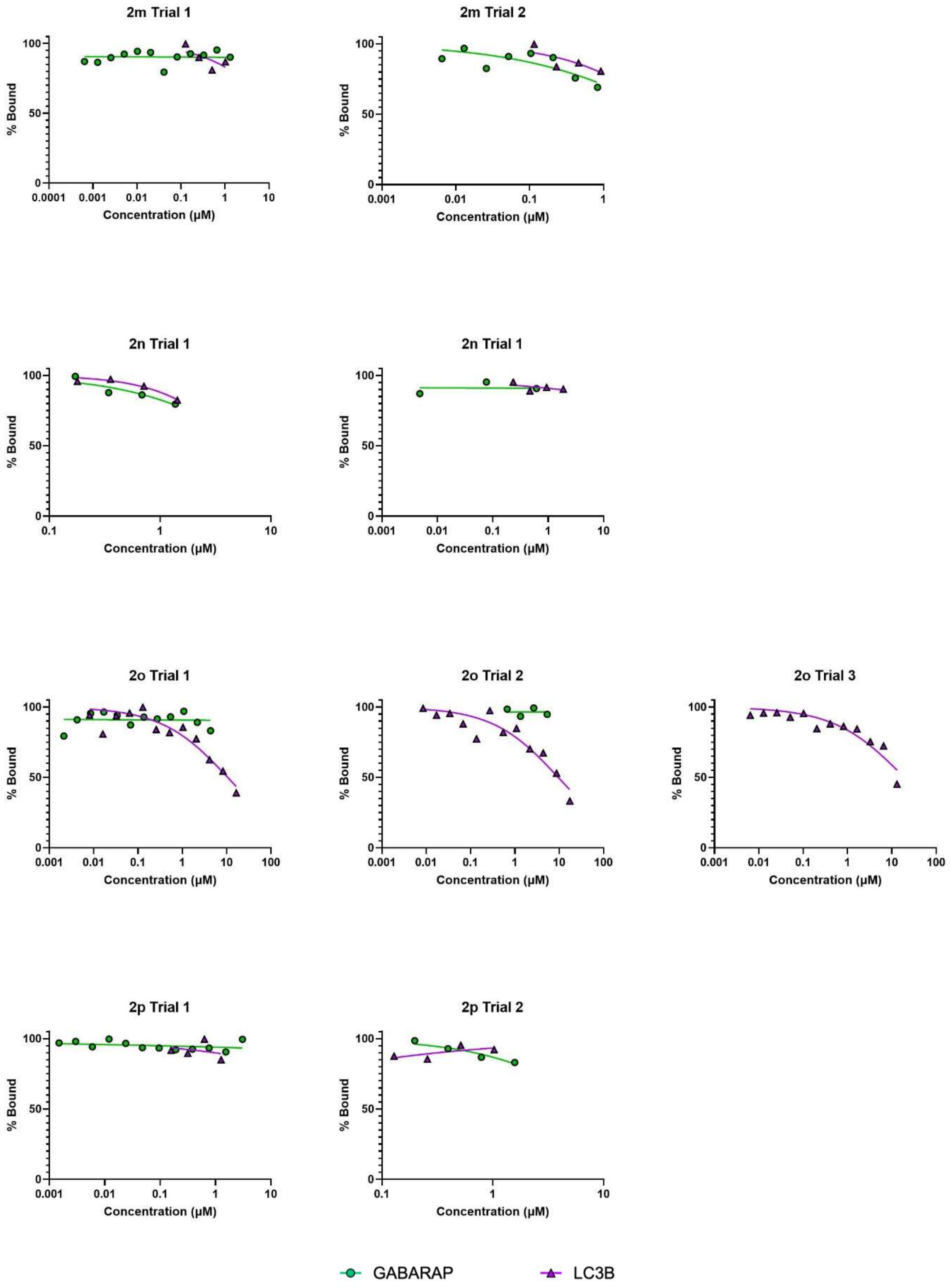
2m Trial 1
2m Trial 2
2n Trial 1
2n Trial 1
2o Trial 1
2o Trial 2
2o Trial 3
2p Trial 1
2p Trial 2
% Bound
Concentration (µM)
GABARAP
LC3B

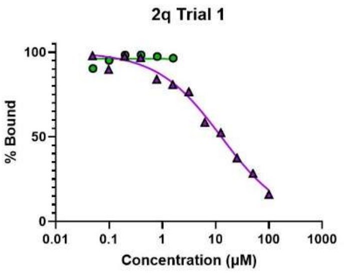
2q Trial 1
% Bound
100
50
0
0.01
0.1
1
10
100
1000
Concentration (μM)

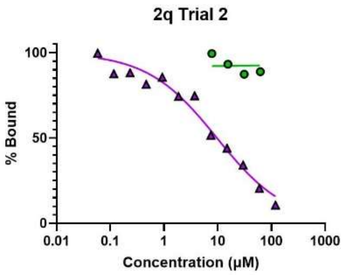
2q Trial 2
% Bound
100
50
0
0.01
0.1
1
10
100
1000
Concentration (μM)

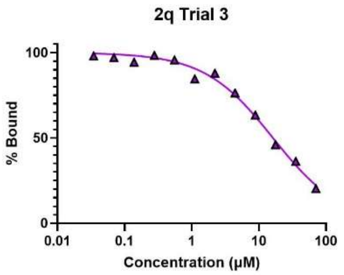
2q Trial 3
% Bound
100
50
0
0.01
0.1
1
10
100
Concentration (μM)

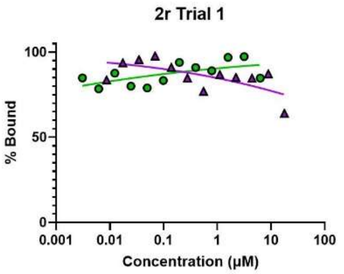
2r Trial 1
% Bound
100
50
0
0.001
0.01
0.1
1
10
100
Concentration (μM)

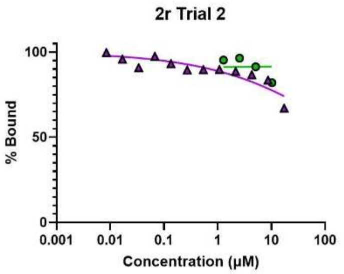
2r Trial 2
% Bound
100
50
0
0.001
0.01
0.1
1
10
100
Concentration (μM)

### 5. NMR Spectra for Novel Compounds

**$^1$H NMR (500 MHz, DMSO-*d*$_6$)** 3-benzylidene-5-iodoindolin-2-one **2a**

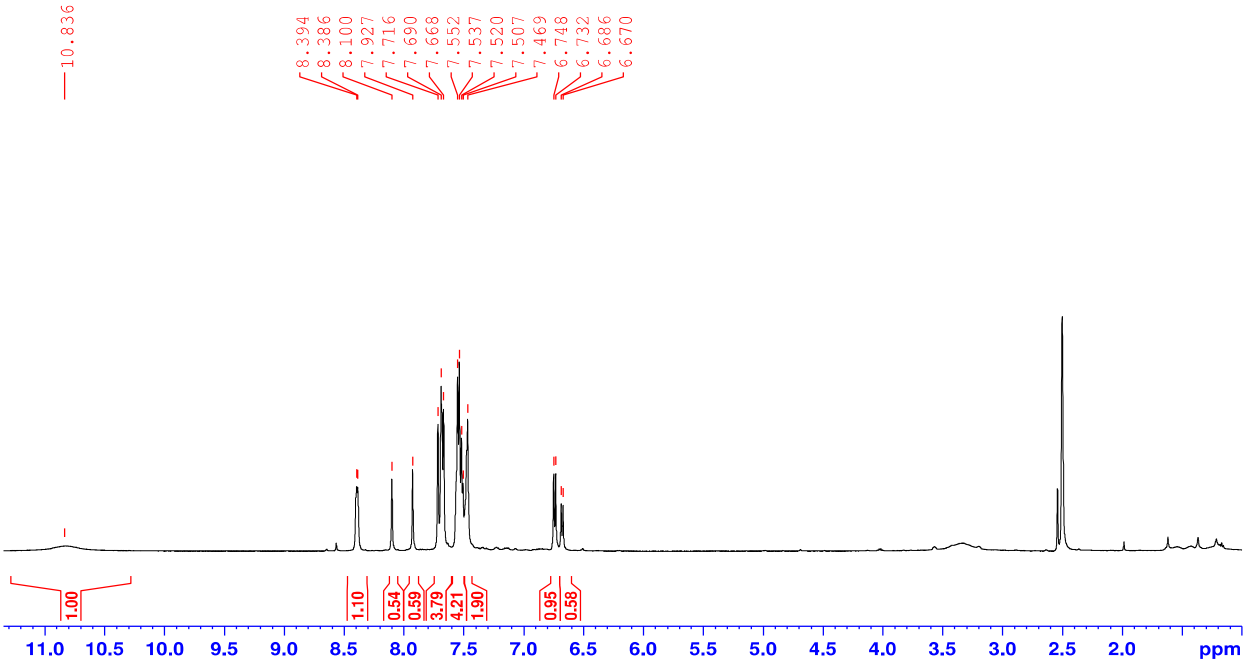


**$^{13}$C NMR (125 MHz, DMSO-*d*$_6$)** 3-benzylidene-5-iodoindolin-2-one **2a**

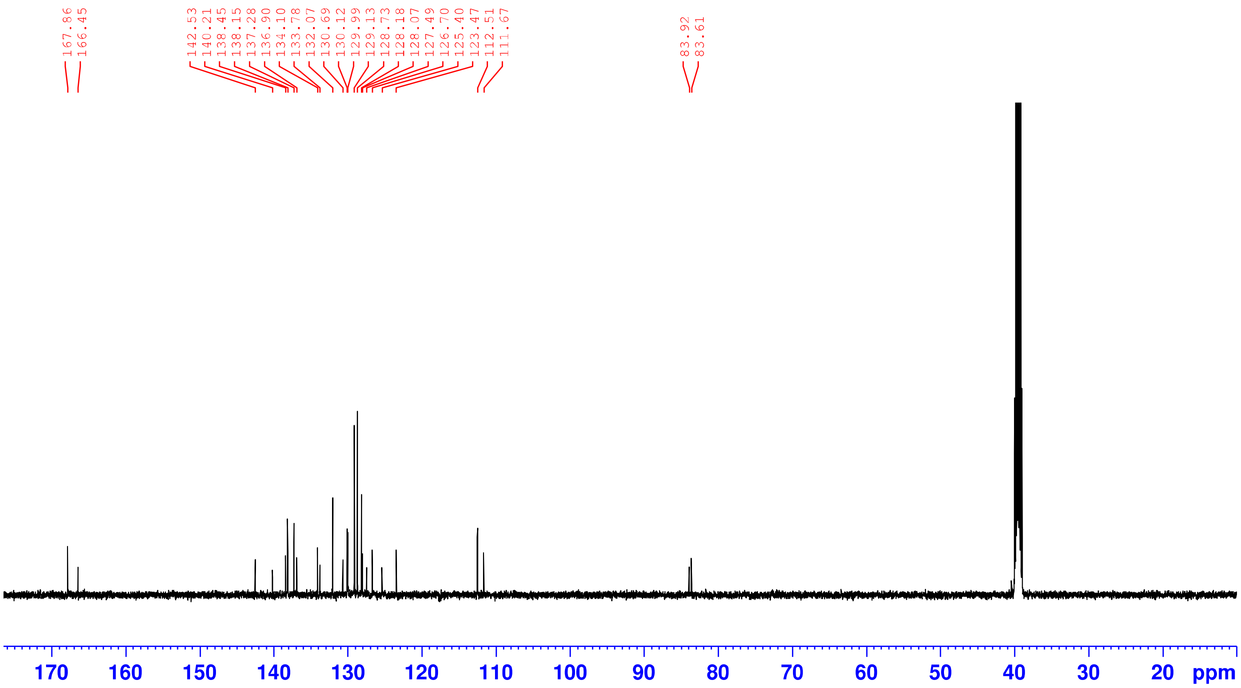

**$^{1}H$ NMR (500 MHz, DMSO-*$d_6$*)** 3-(4-hydroxybenzylidene)-5-iodoindolin-2-one **2b**

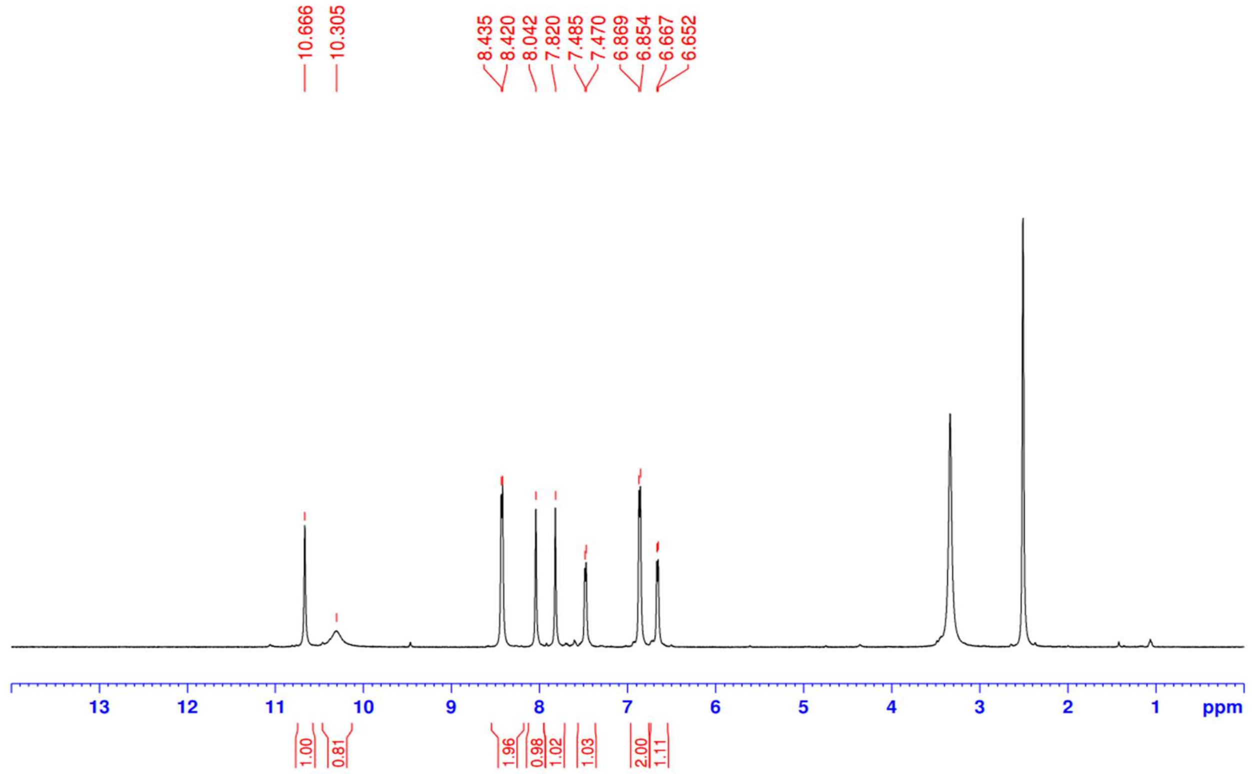


**$^{13}C$ NMR (125 MHz, DMSO-*$d_6$*)** 3-(4-hydroxybenzylidene)-5-iodoindolin-2-one **2b**

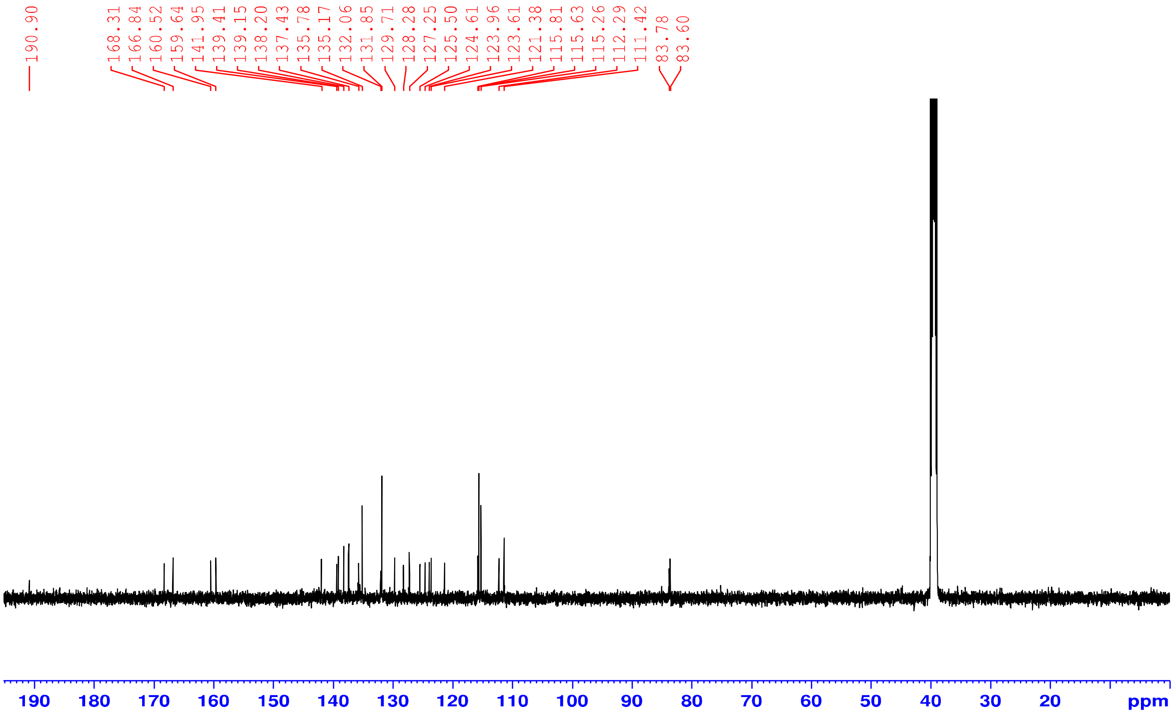

**$^{1}$H NMR (500 MHz, DMSO-*d*$_6$)** 3-(3,5-dibromobenzylidene)-5-iodoindolin-2-one **2c**

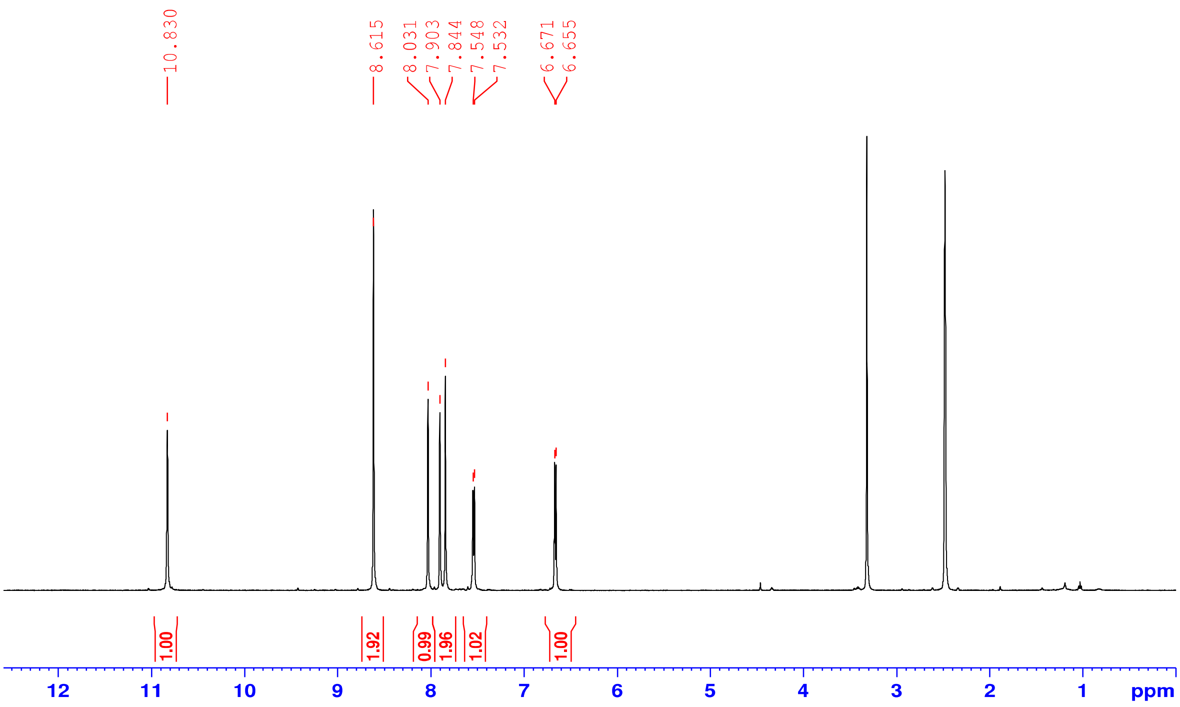


**$^{13}$C NMR (125 MHz, DMSO-*d*$_6$)** 3-(3,5-dibromobenzylidene)-5-iodoindolin-2-one **2c**

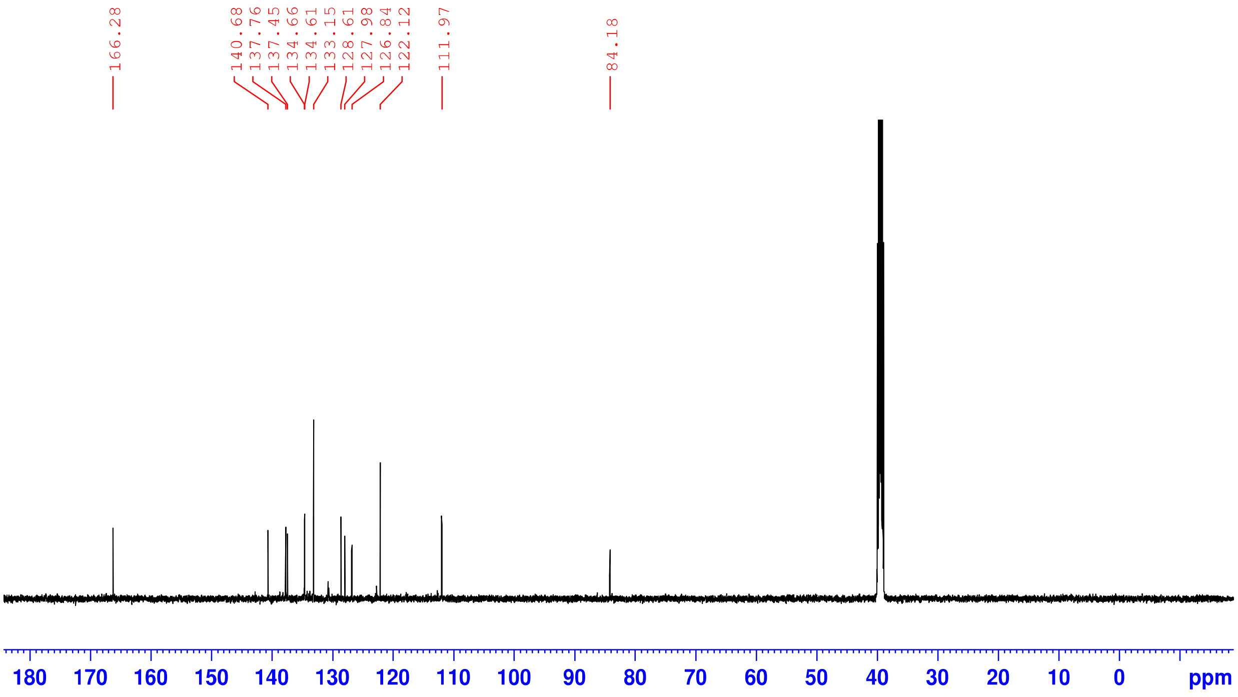

**$^1$H NMR (500 MHz, DMSO-*d$_6$*)** 3-(3,5-dibromo-2-hydroxybenzylidene)-5-iodoindolin-2-one **2d**

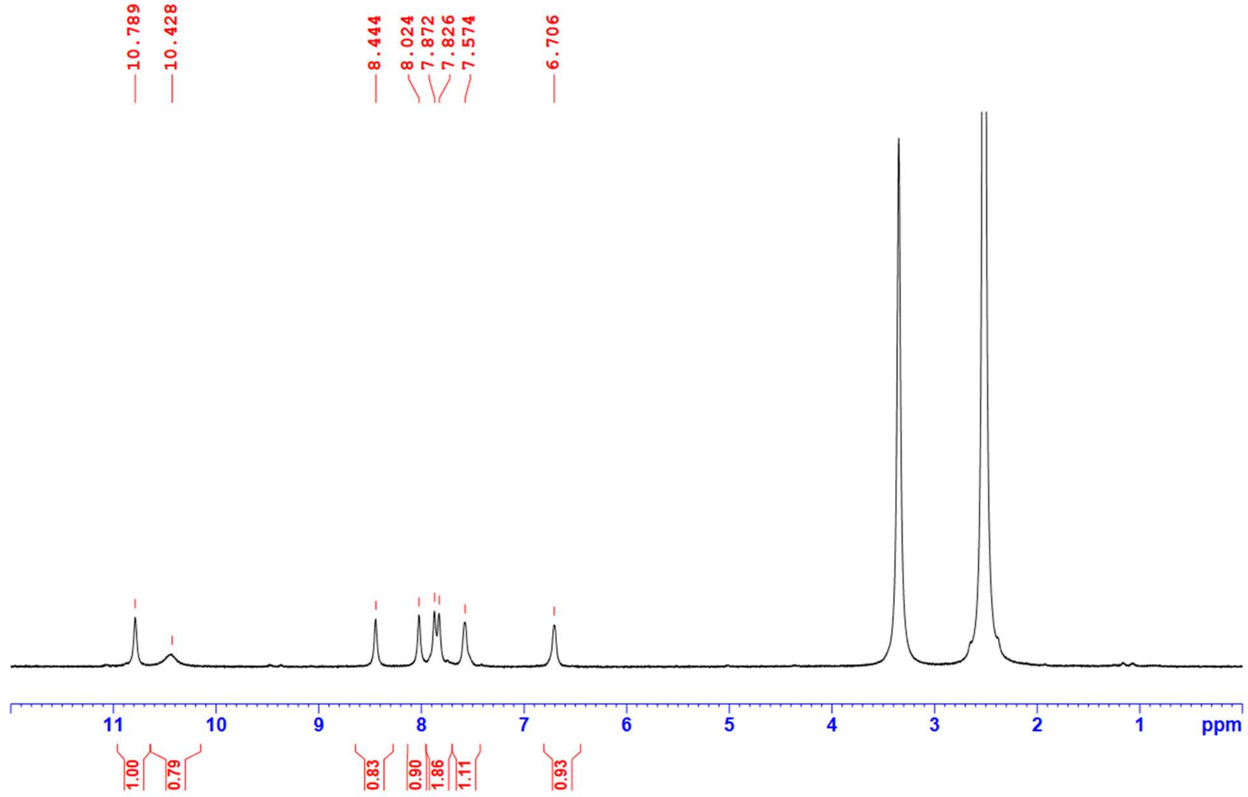


**$^{13}$C NMR (125 MHz, DMSO-*d$_6$*)** 3-(3,5-dibromo-2-hydroxybenzylidene)-5-iodoindolin-2-one **2d**

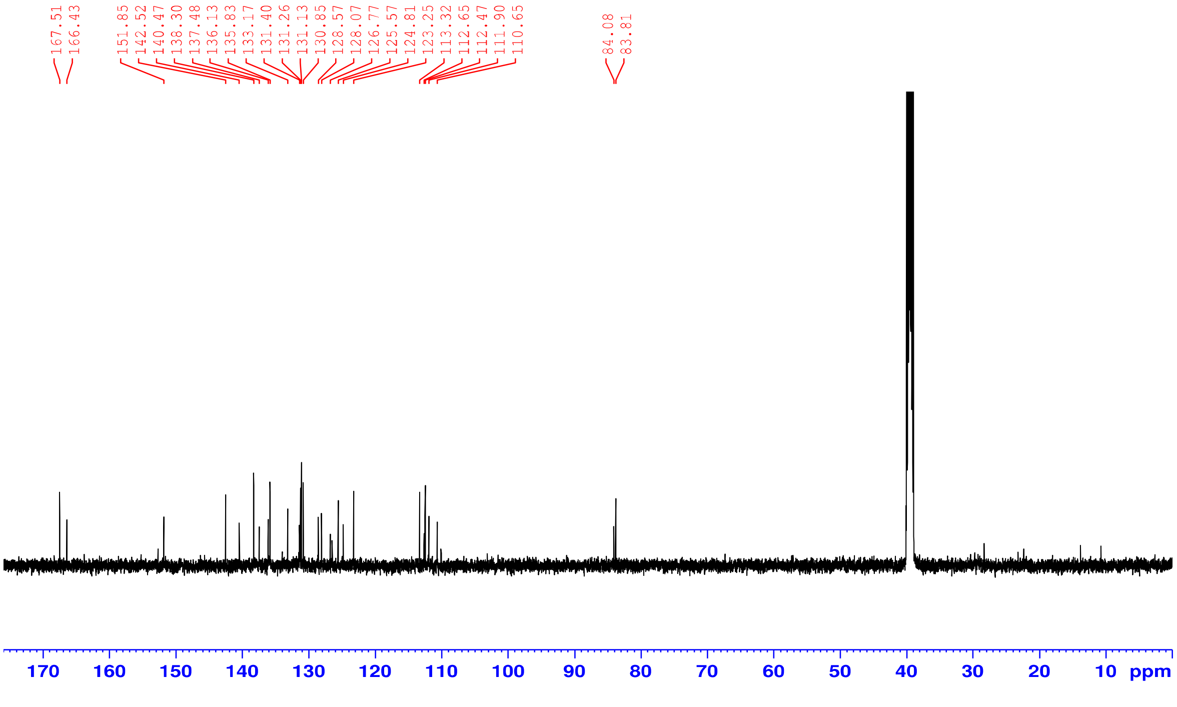

**$^1$H NMR (500 MHz, DMSO-*$d_6$*)** 5-iodo-3-(4-methoxybenzylidene)indolin-2-one **2e**

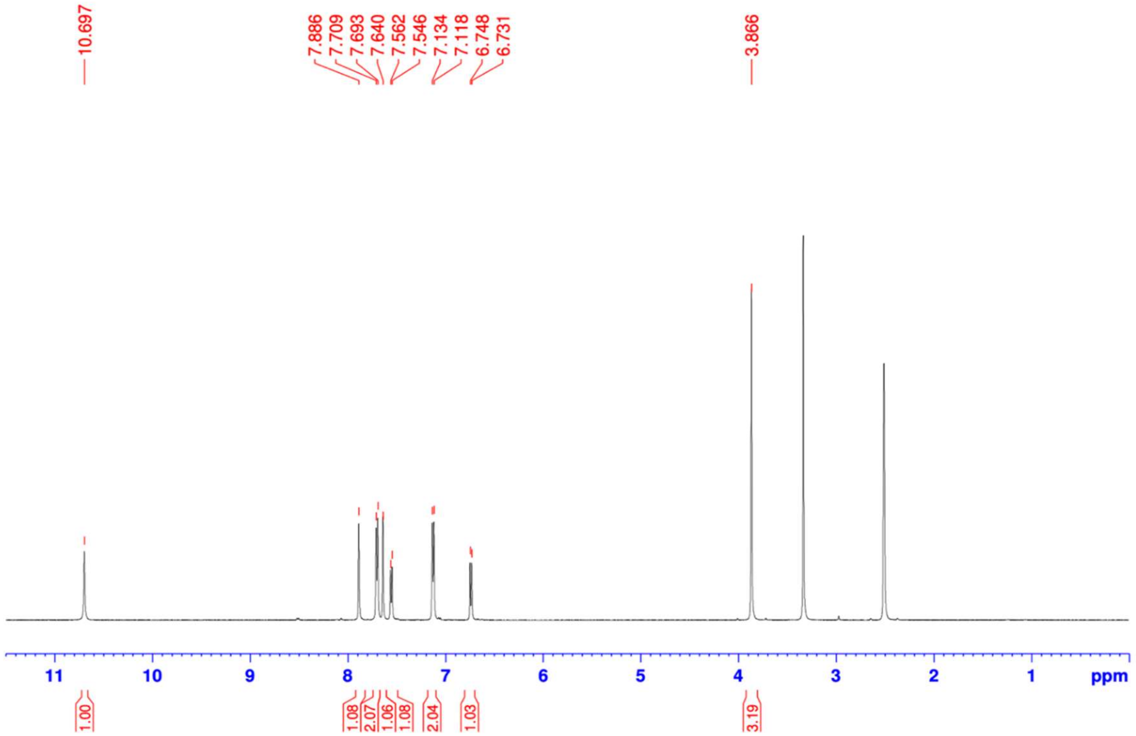


**$^{13}$C NMR (125 MHz, DMSO-*$d_6$*)** 5-iodo-3-(4-methoxybenzylidene)indolin-2-one **2e**

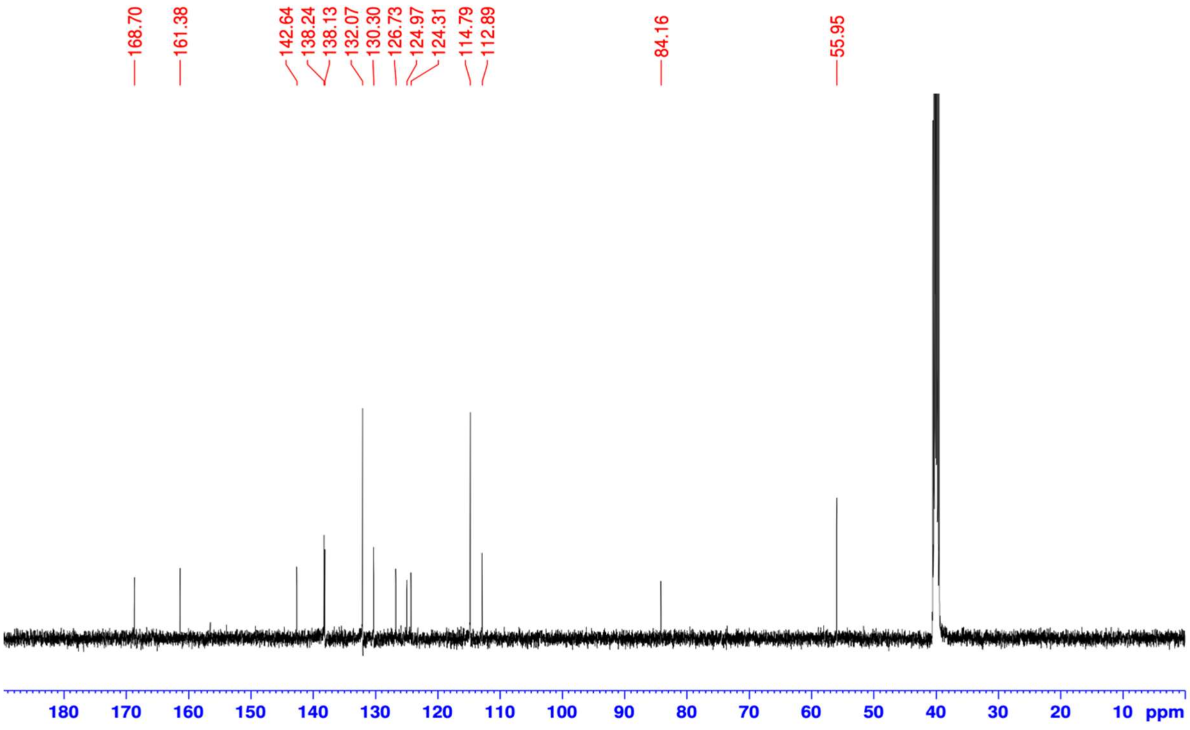

**[1]H NMR (500 MHz, DMSO-*d6*)** 4-((5-iodo-2-oxoindolin-3-ylidene)methyl)benzoic acid **2f**

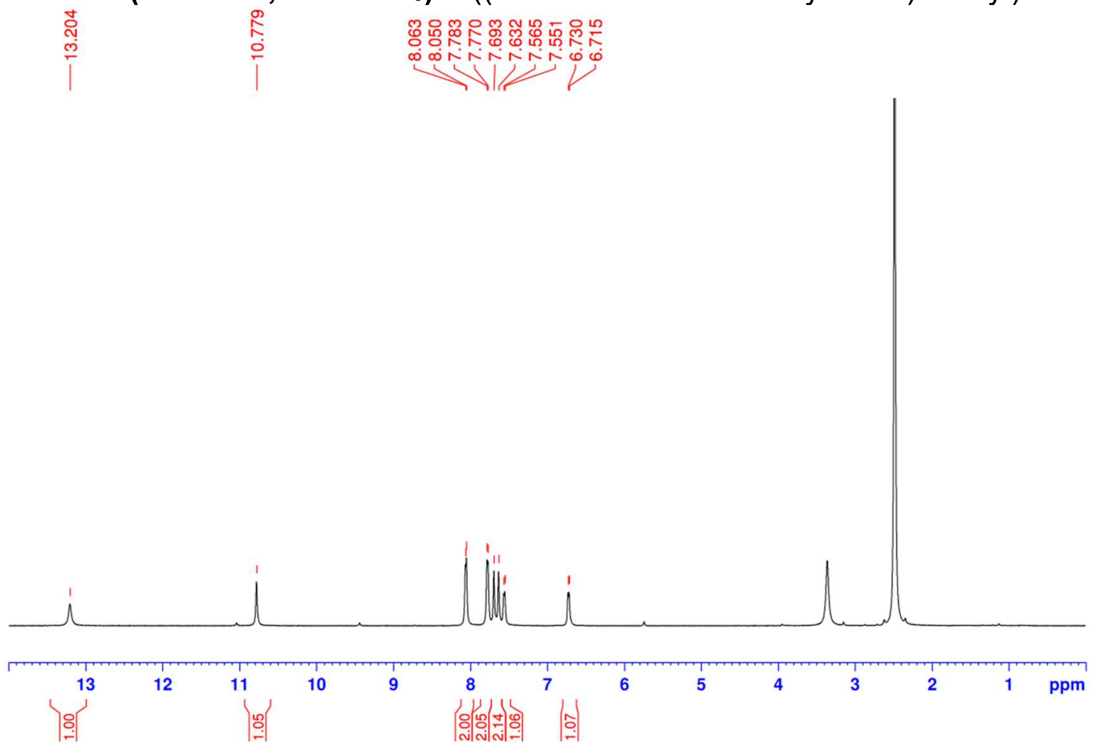


**[13]C NMR (125 MHz, DMSO-*d6*)** 4-((5-iodo-2-oxoindolin-3-ylidene)methyl)benzoic acid **2f**

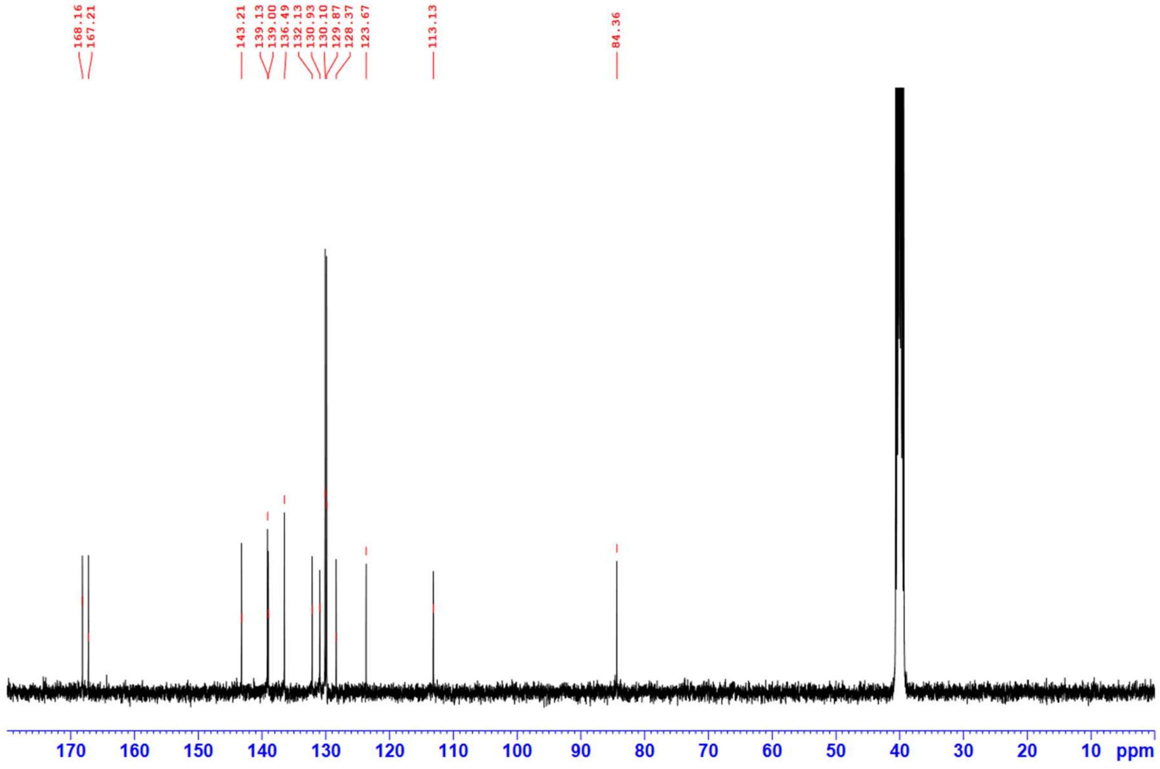

**$^1$H NMR (500 MHz, DMSO-$d_6$)** 3-(3,5-dibromo-4-hydroxybenzylidene)indolin-2-one **2g**

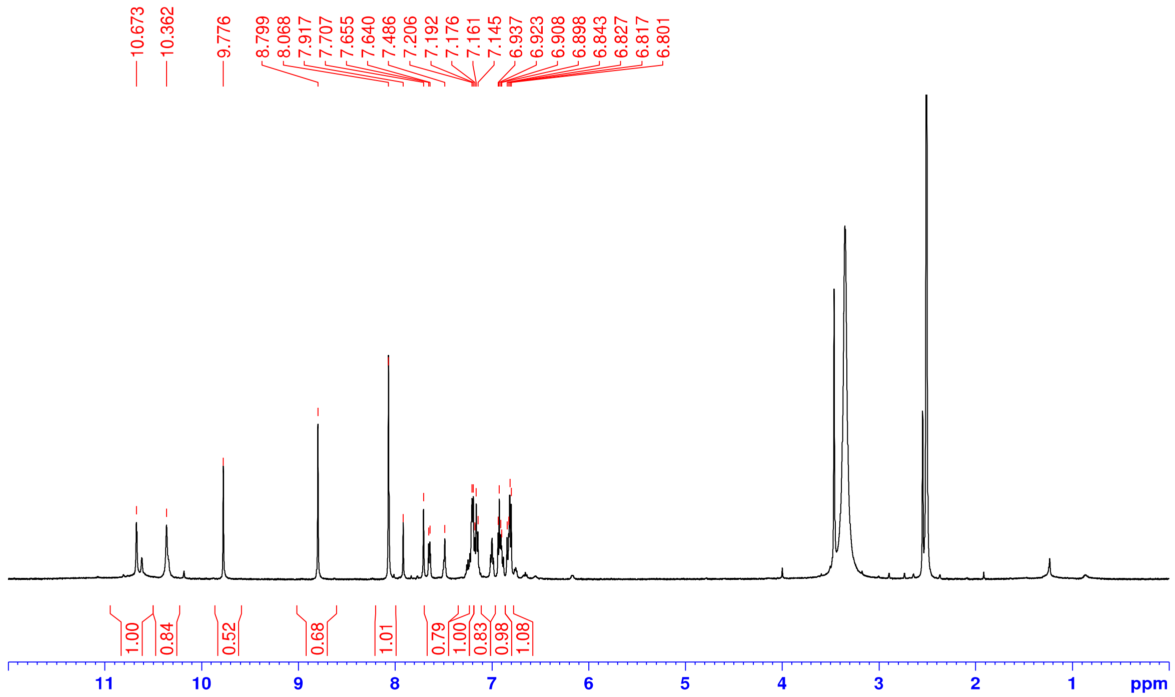


**$^{13}$C NMR (125 MHz, DMSO-$d_6$)** 3-(3,5-dibromo-4-hydroxybenzylidene)indolin-2-one **2g**

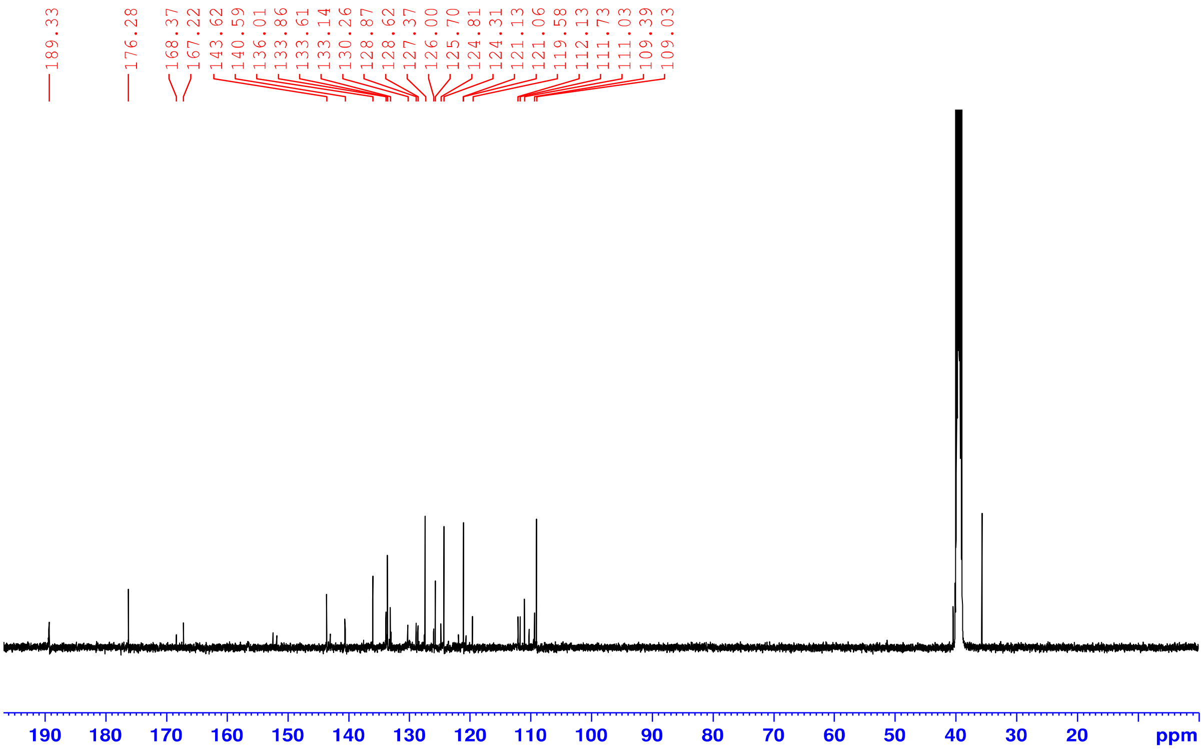

**$^{1}$H NMR (500 MHz, DMSO-*d₆*)** 3-(3,5-dibromo-4-hydroxybenzylidene)-6-iodoindolin-2-one **2h**

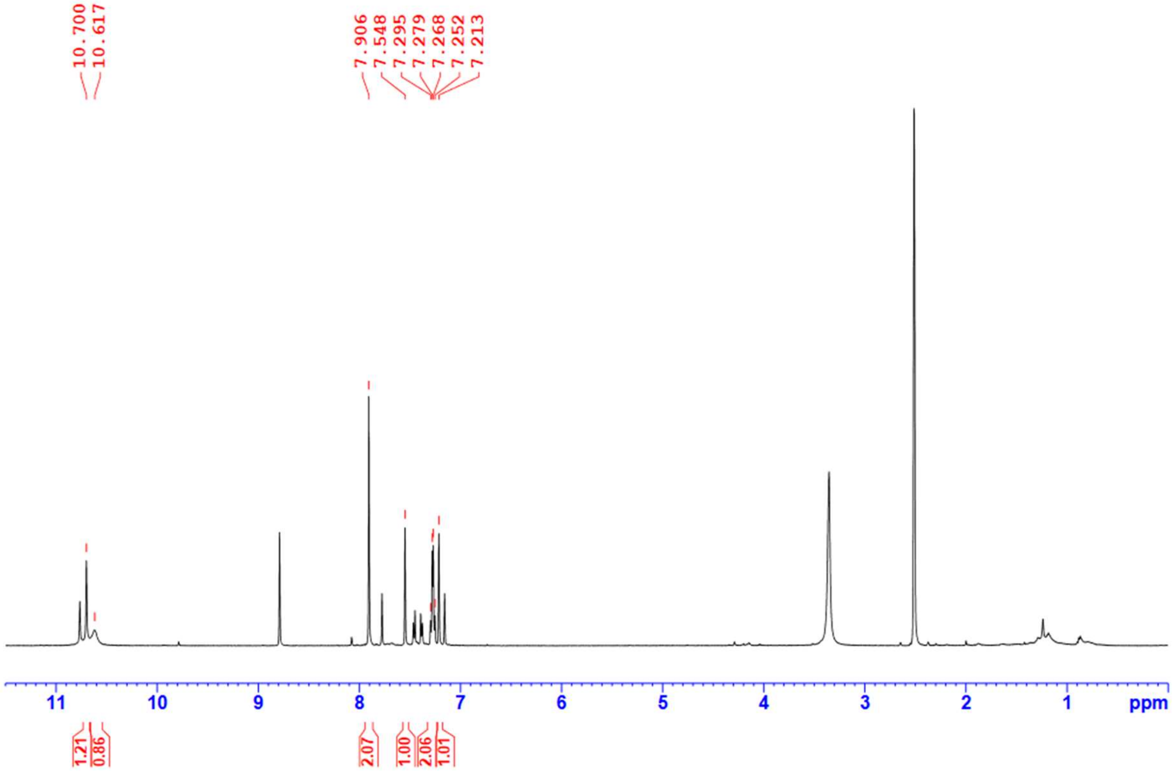


**$^{13}$C NMR (125 MHz, DMSO-*d₆*)** 3-(3,5-dibromo-4-hydroxybenzylidene)-6-iodoindolin-2-one **2h**

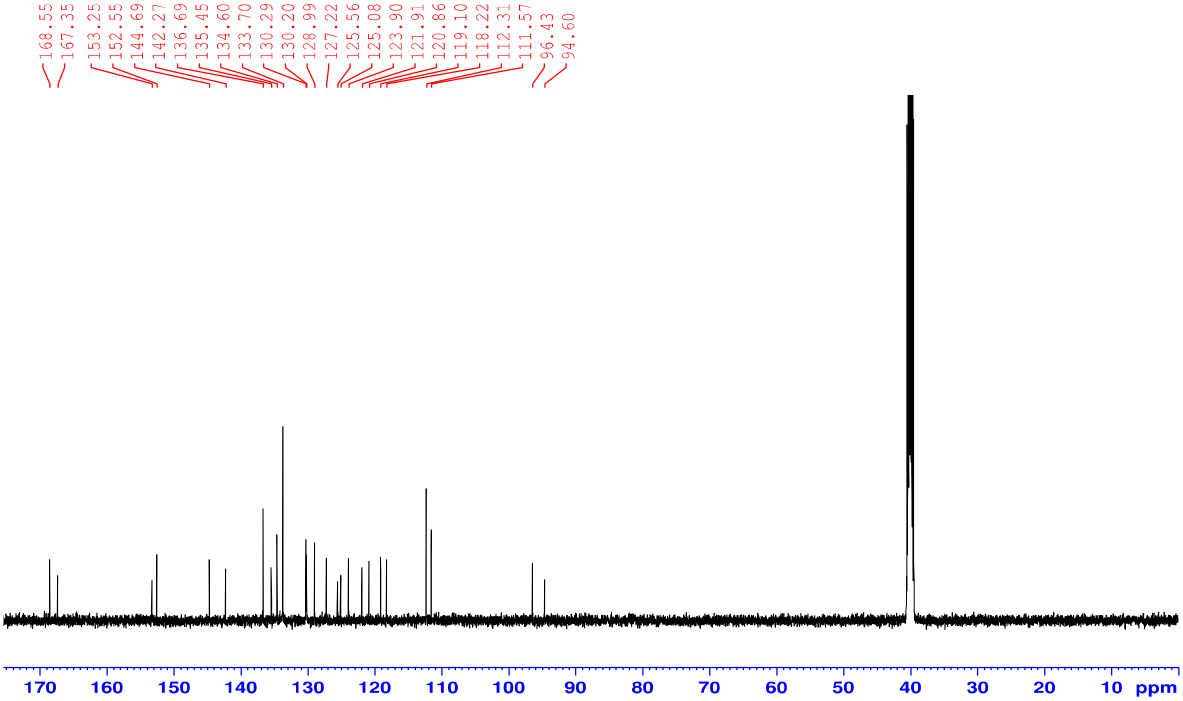

**$^{1}$H NMR (500 MHz, DMSO-*d$_6$*)** 5-bromo-3-(3,5-dibromo-4-hydroxybenzylidene)indolin-2-one **2i**

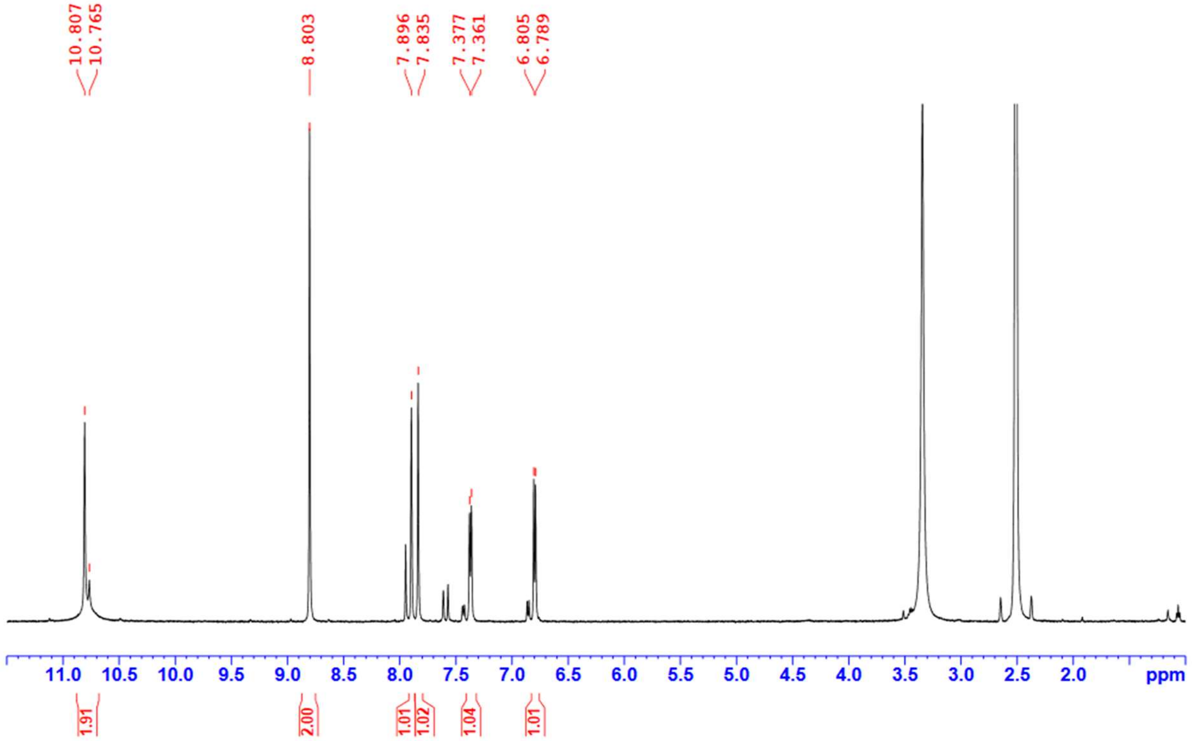


**$^{13}$C NMR (125 MHz, DMSO-*d$_6$*)** 5-bromo-3-(3,5-dibromo-4-hydroxybenzylidene)indolin-2-one **2i**

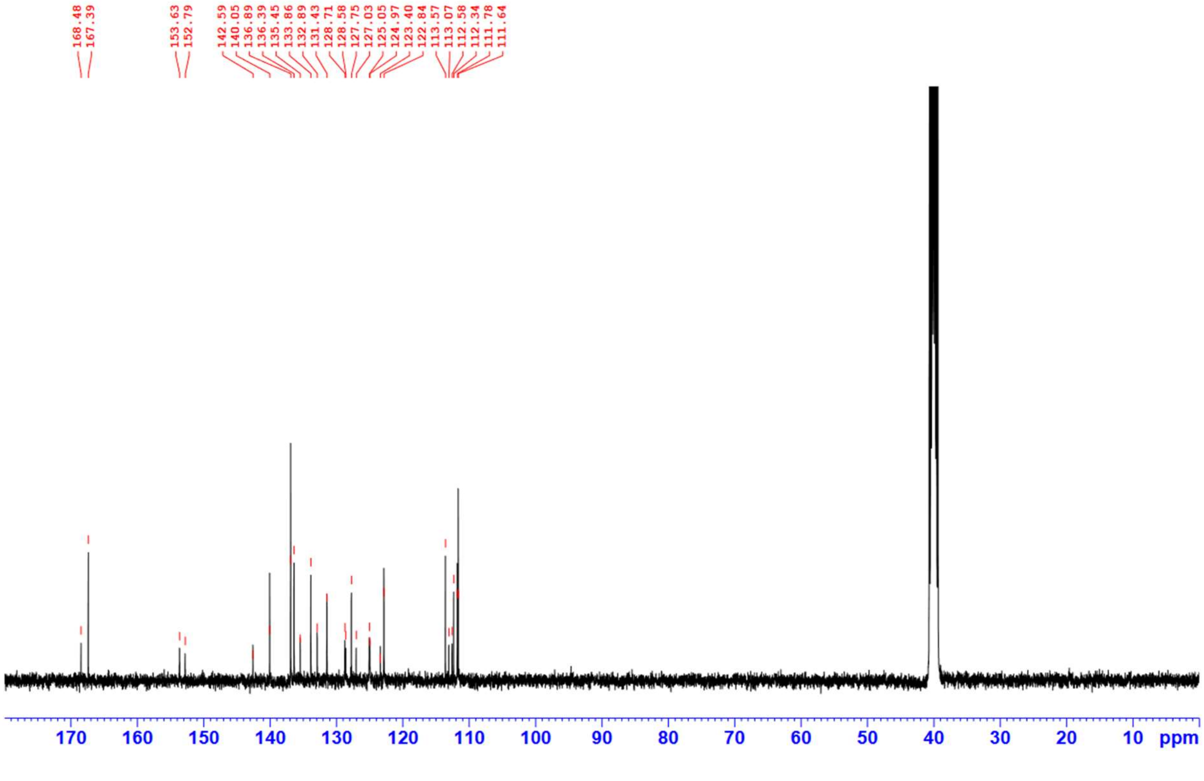

**$^{1}$H NMR (500 MHz, DMSO-*$d_6$*)** 3-(3,5-dibromo-4-hydroxybenzylidene)-6-methylindolin-2-one **2j**

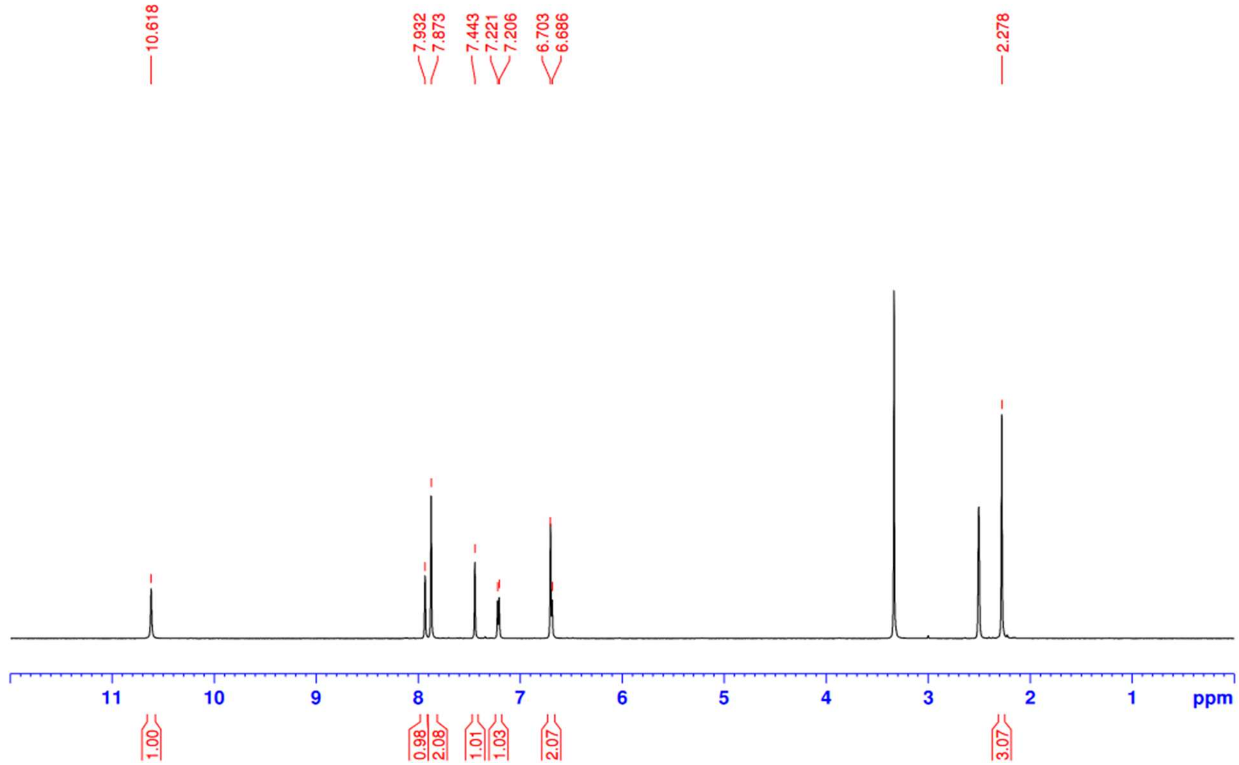

**$^{13}$C NMR (125 MHz, DMSO-*$d_6$*)** 3-(3,5-dibromo-4-hydroxybenzylidene)-6-methylindolin-2-one **2j**

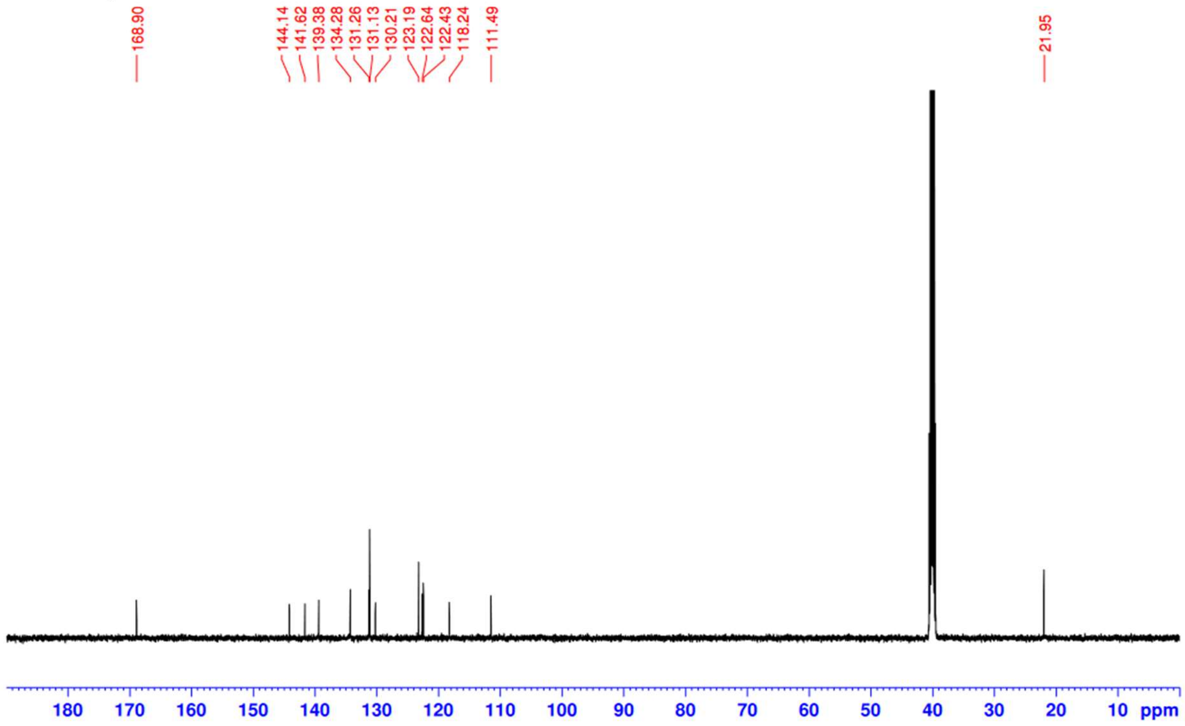

**¹H NMR (500 MHz, DMSO-*d₆*)** methyl 3-(3,5-dibromo-4-hydroxybenzylidene)-2-oxoindoline-6-carboxylate **2k**

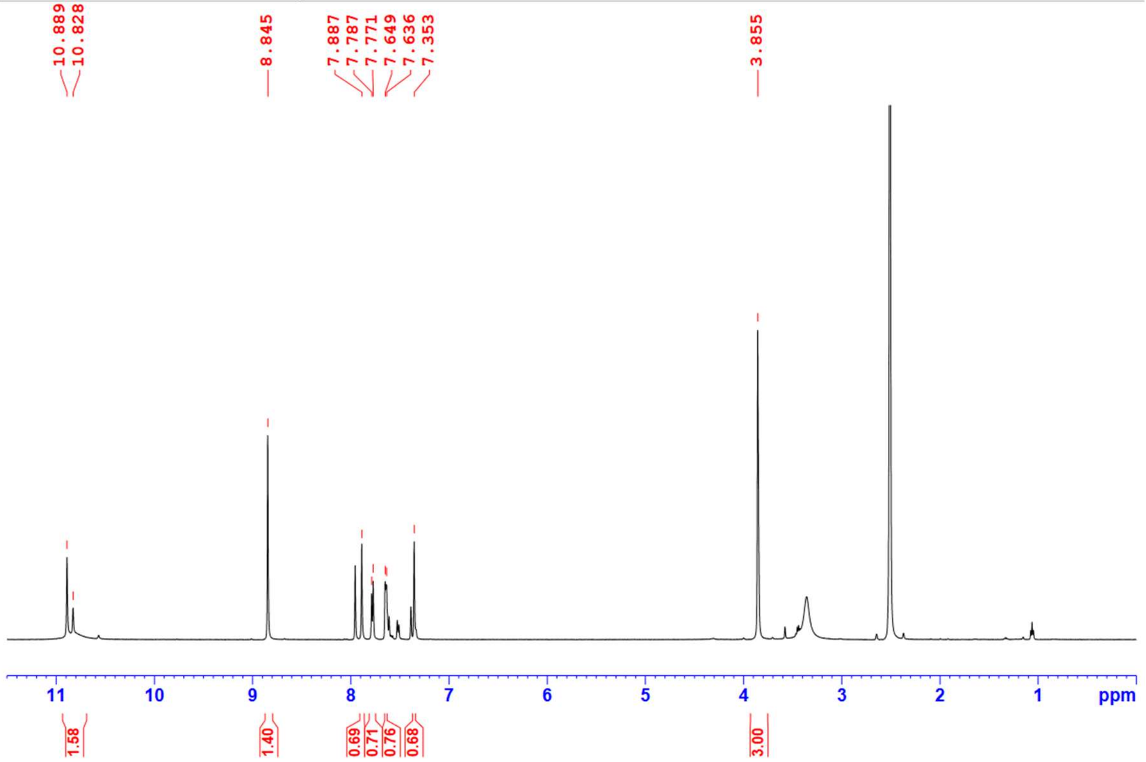


**¹³C NMR (125 MHz, DMSO-*d₆*)** methyl 3-(3,5-dibromo-4-hydroxybenzylidene)-2-oxoindoline-6-carboxylate **2k**

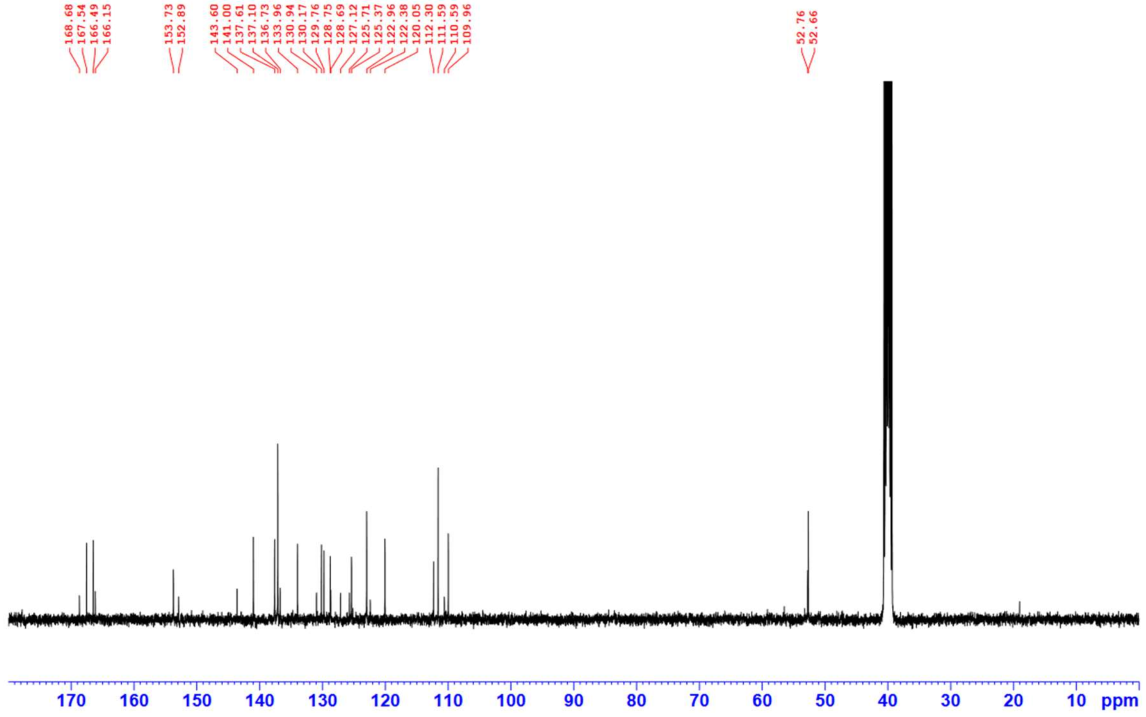

**$^{1}$H NMR (500 MHz, DMSO-$d_6$)** 5,6-dichloro-3-(3,5-dibromo-4-hydroxybenzylidene)indolin-2-one **2l**

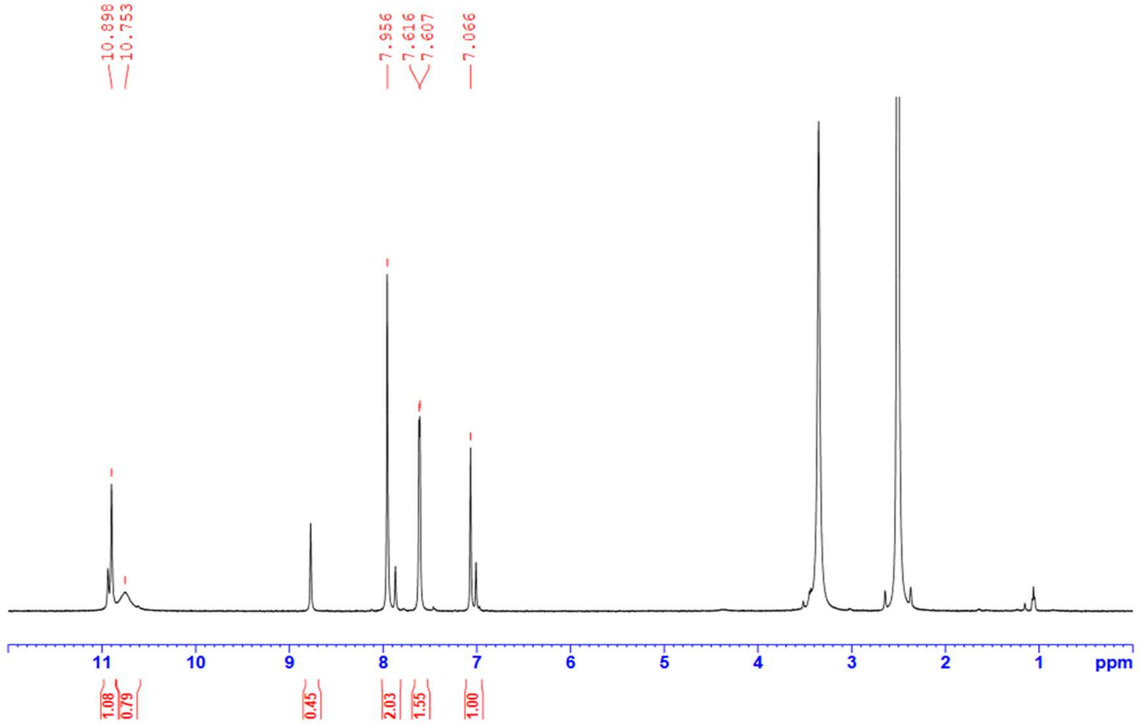


**$^{13}$C NMR (125 MHz, DMSO-$d_6$)** 5,6-dichloro-3-(3,5-dibromo-4-hydroxybenzylidene)indolin-2-one **2l**

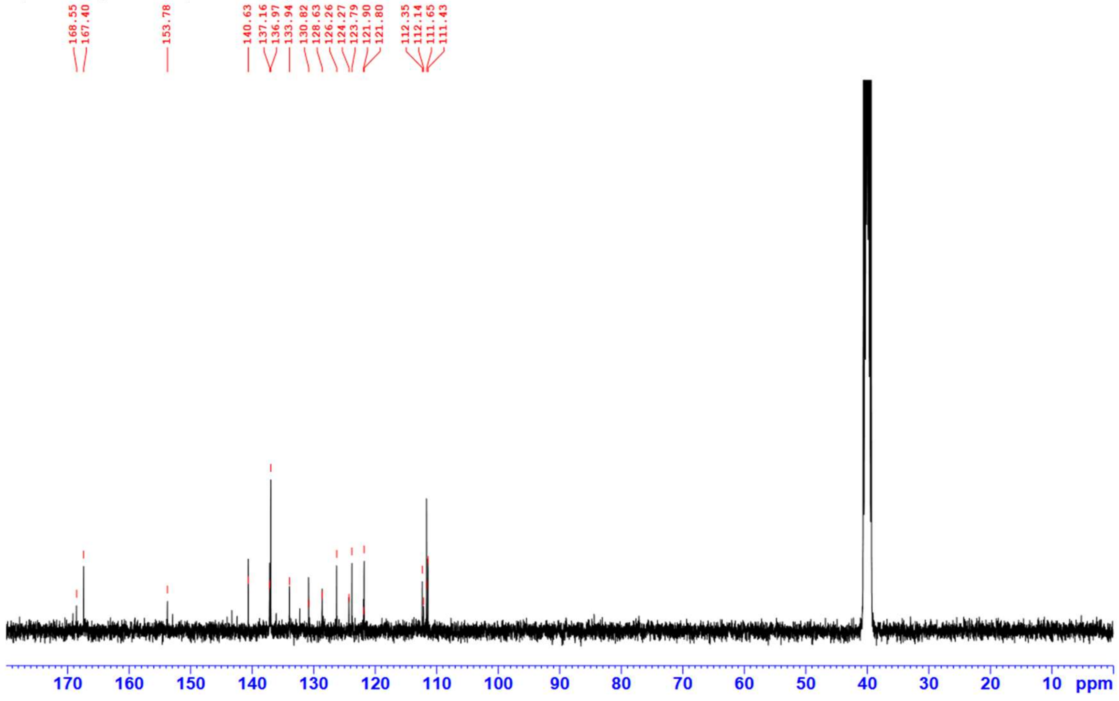

**$^{1}H$ NMR (500 MHz, DMSO-*$d_6$*)** 5-iodo-3-((4'-(trifluoromethyl)-[1,1'-biphenyl]-4-yl)methylene)indolin-2-one **2m**

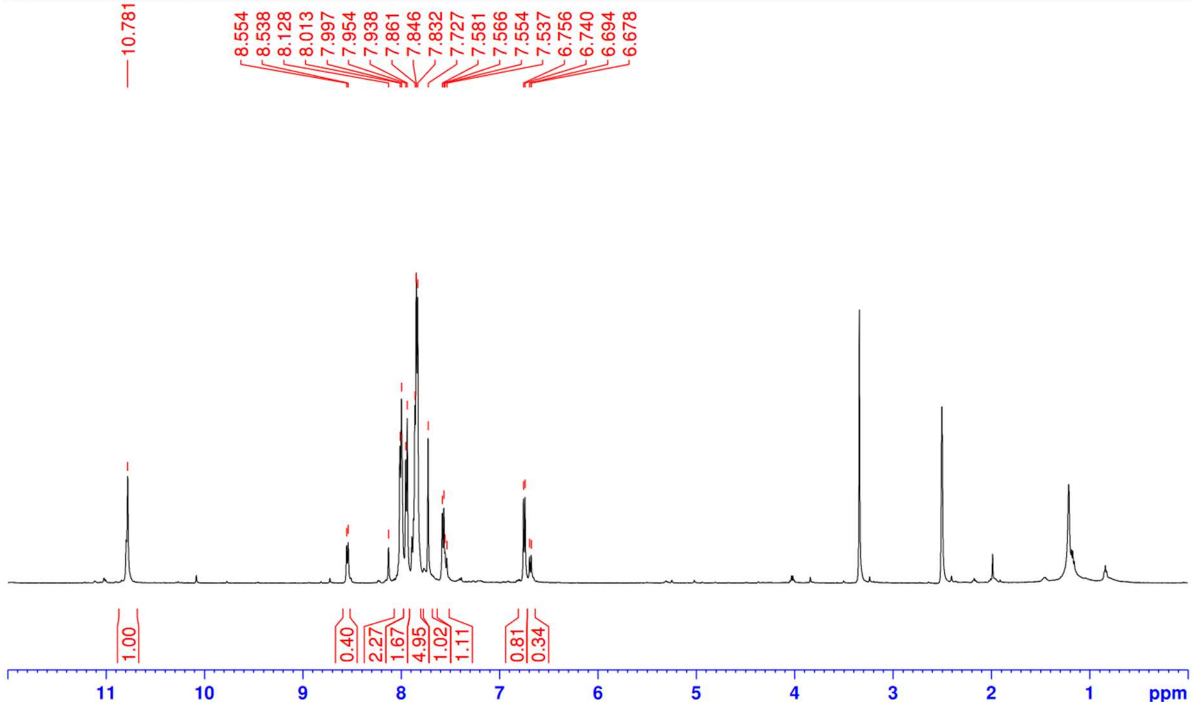


**$^{13}C$ NMR (125 MHz, DMSO-*$d_6$*)** 5-iodo-3-((4'-(trifluoromethyl)-[1,1'-biphenyl]-4-yl)methylene)indolin-2-one **2m**

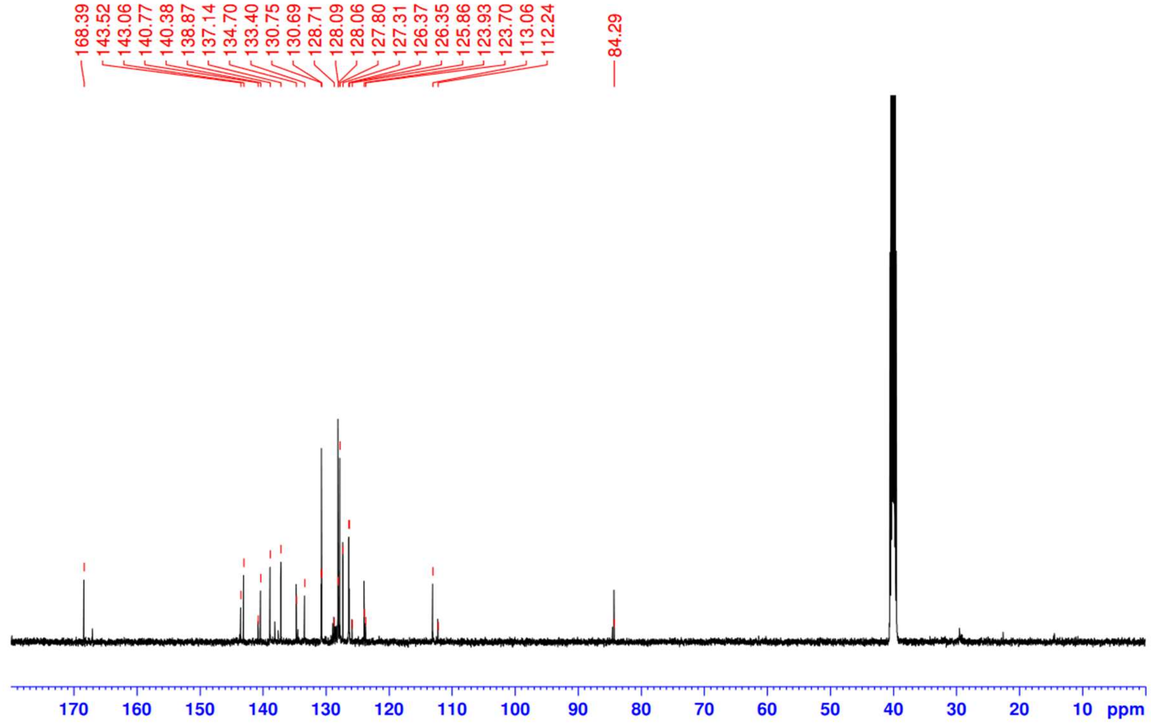

**$^{1}H$ NMR (500 MHz, DMSO-*$d_6$*)** 5-iodo-3-(thiazol-5-ylmethylene)indolin-2-one **2n**

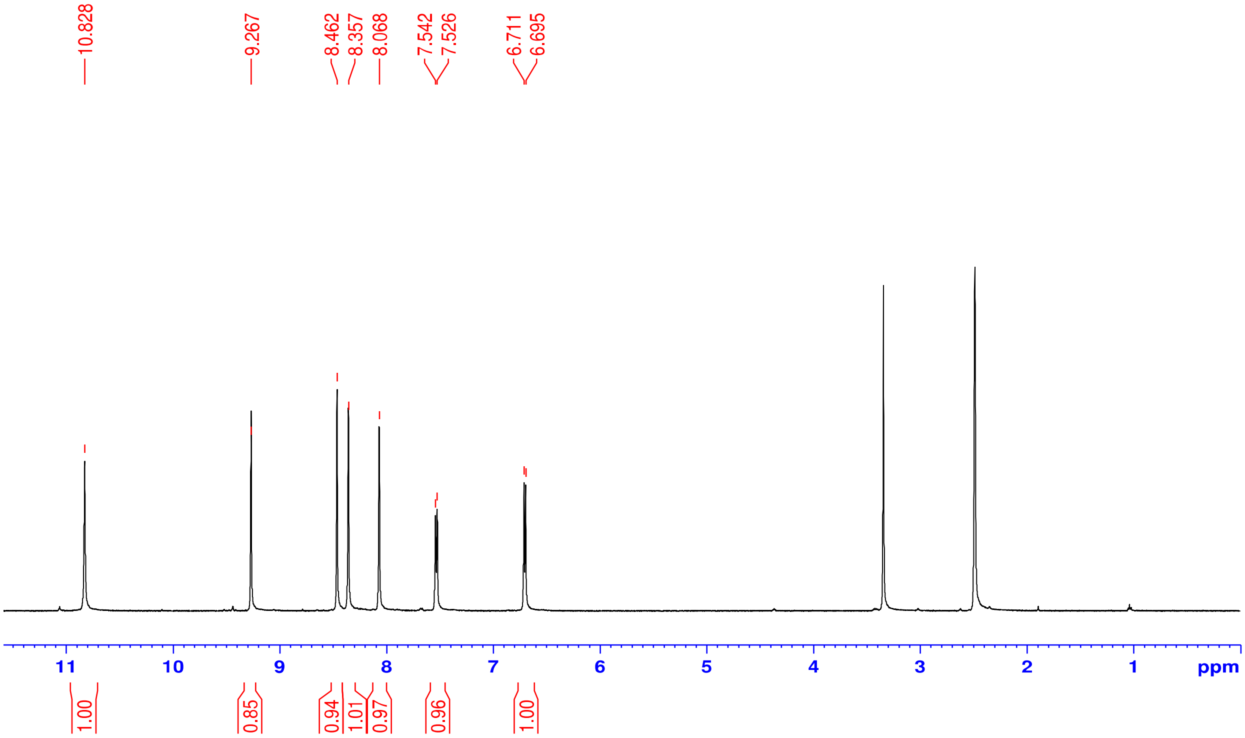


**$^{13}C$ NMR (125 MHz, DMSO-*$d_6$*)** 5-iodo-3-(thiazol-5-ylmethylene)indolin-2-one **2n**

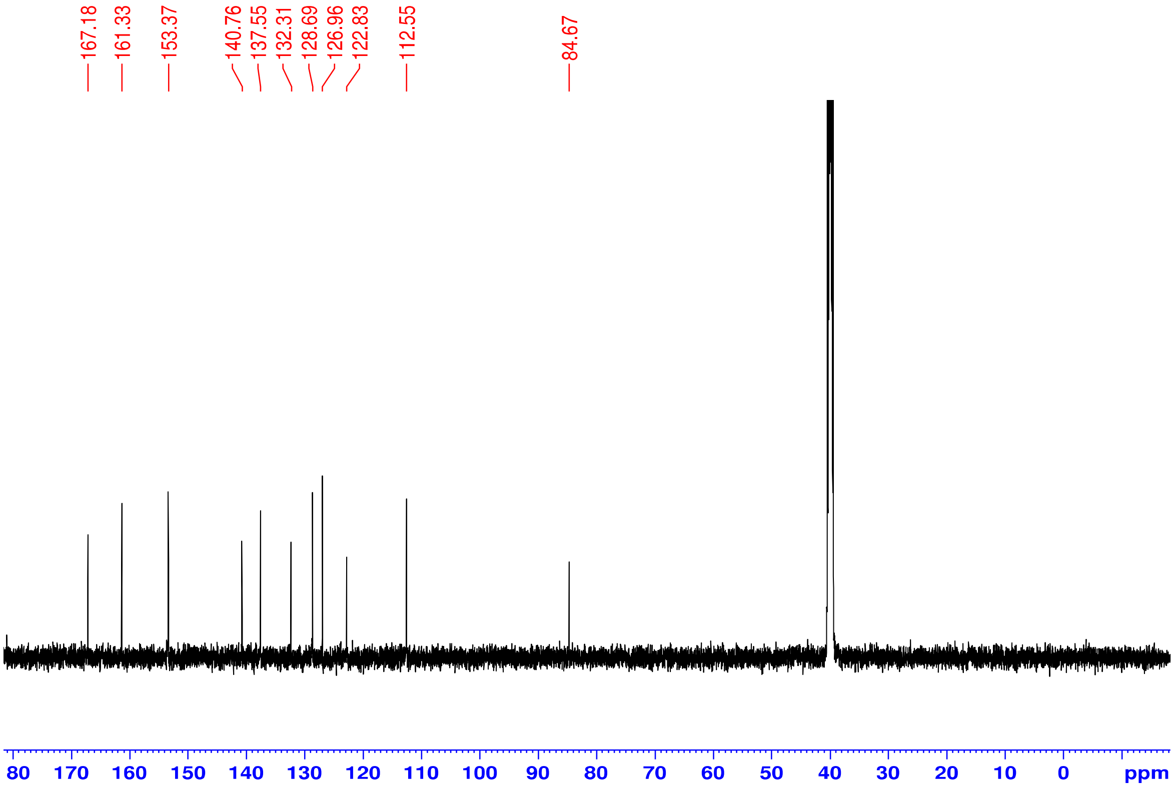

**[1]H NMR (500 MHz, DMSO-$d_6$)** 5-iodo-3-((4-methylthiazol-5-yl)methylene)indolin-2-one **2o**

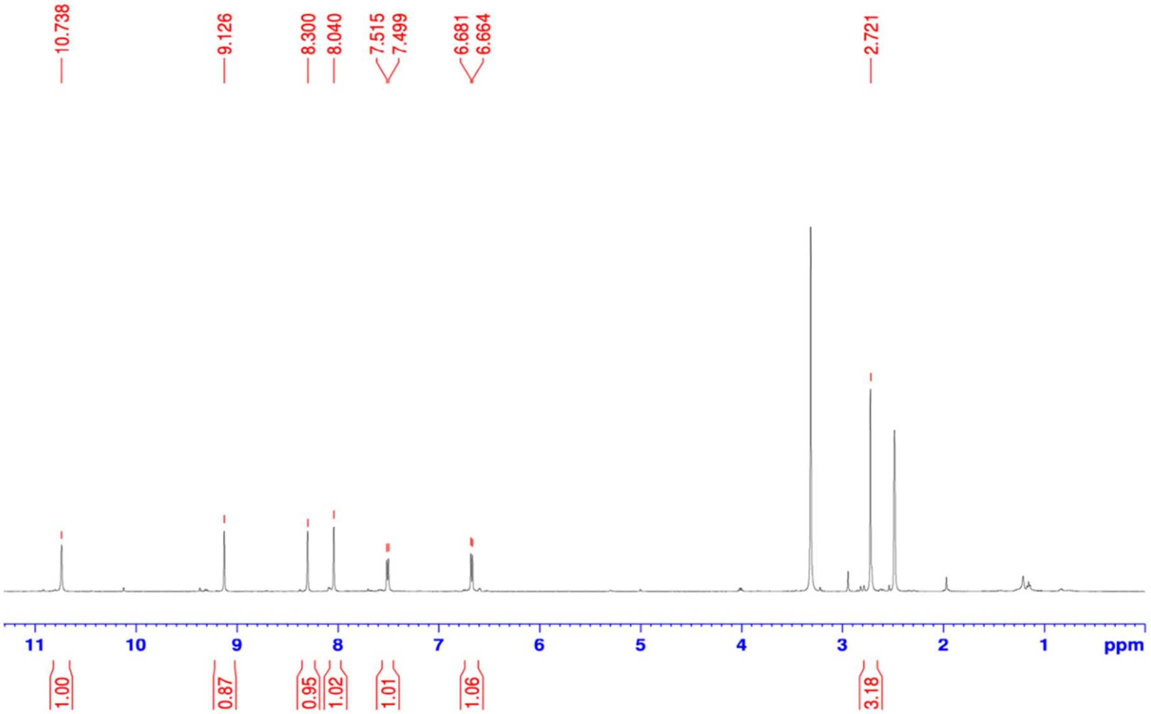


**[13]C NMR (125 MHz, DMSO-$d_6$)** 5-iodo-3-((4-methylthiazol-5-yl)methylene)indolin-2-one **2o**

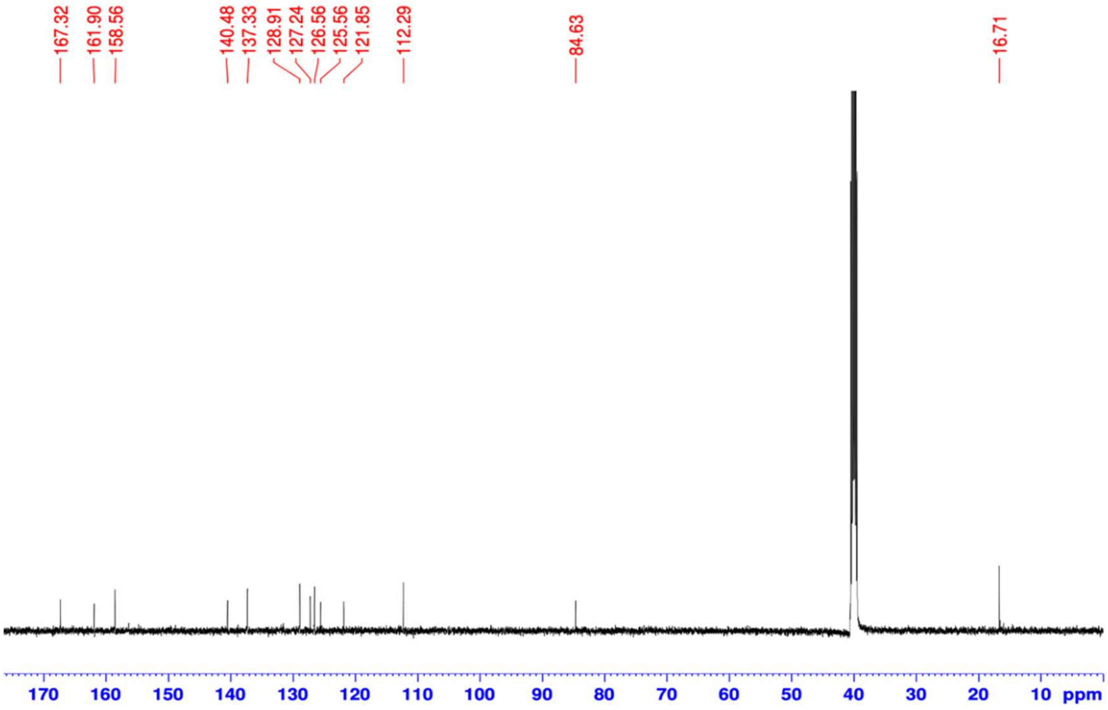

[1]H NMR (500 MHz, DMSO-$d_6$) 3-((1H-pyrrol-2-yl)methylene)-5-iodoindolin-2-one **2p**

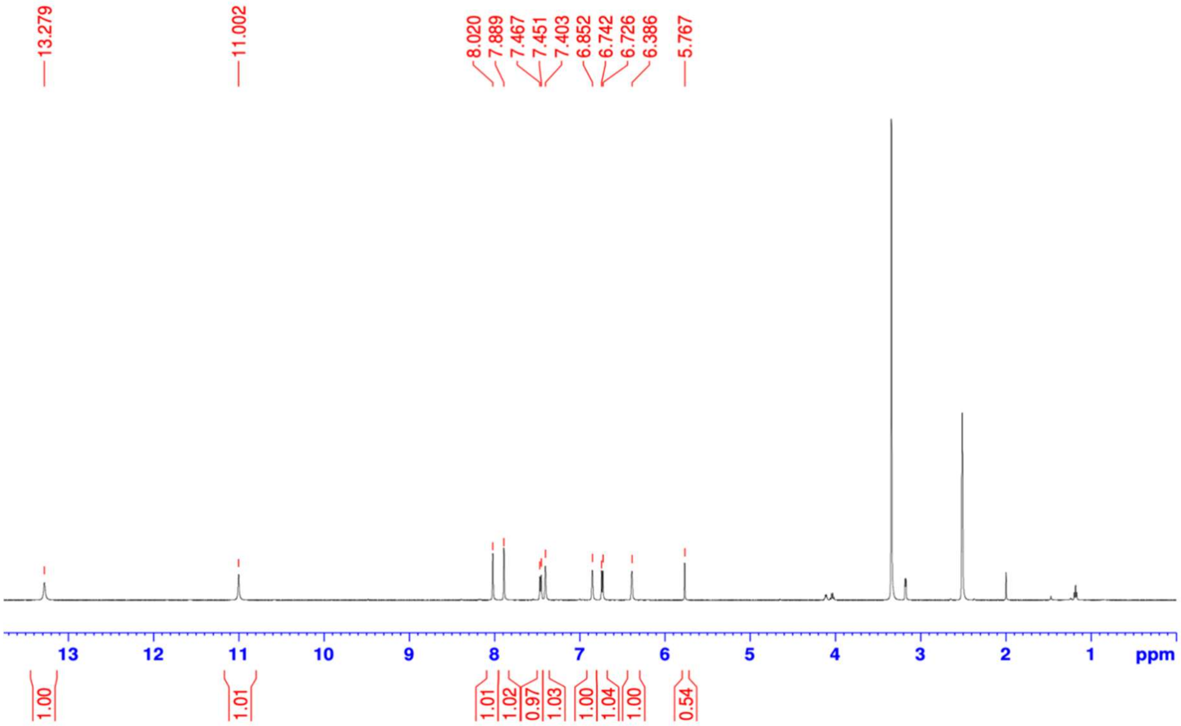


[13]C NMR (125 MHz, DMSO-$d_6$) 3-((1H-pyrrol-2-yl)methylene)-5-iodoindolin-2-one **2p**

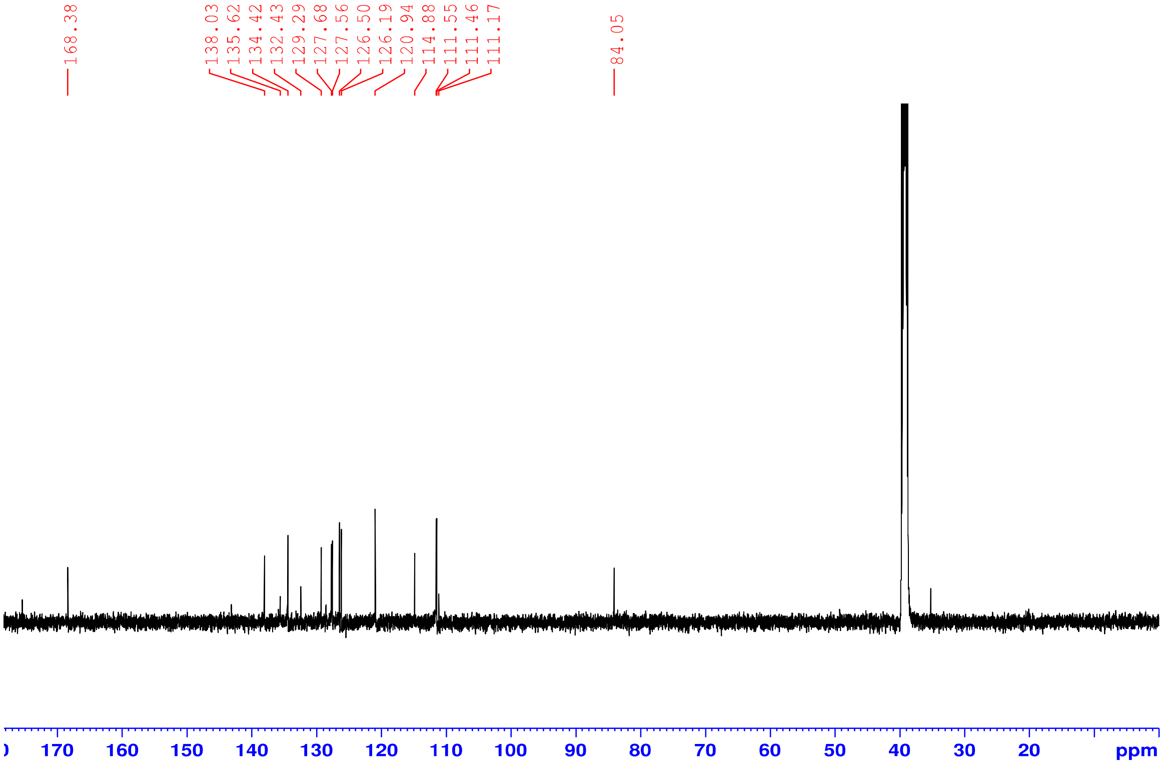

**$^1$H NMR (500 MHz, DMSO-*d₆*)** 3-((1H-imidazol-5-yl)methylene)-5-iodoindolin-2-one **2q**

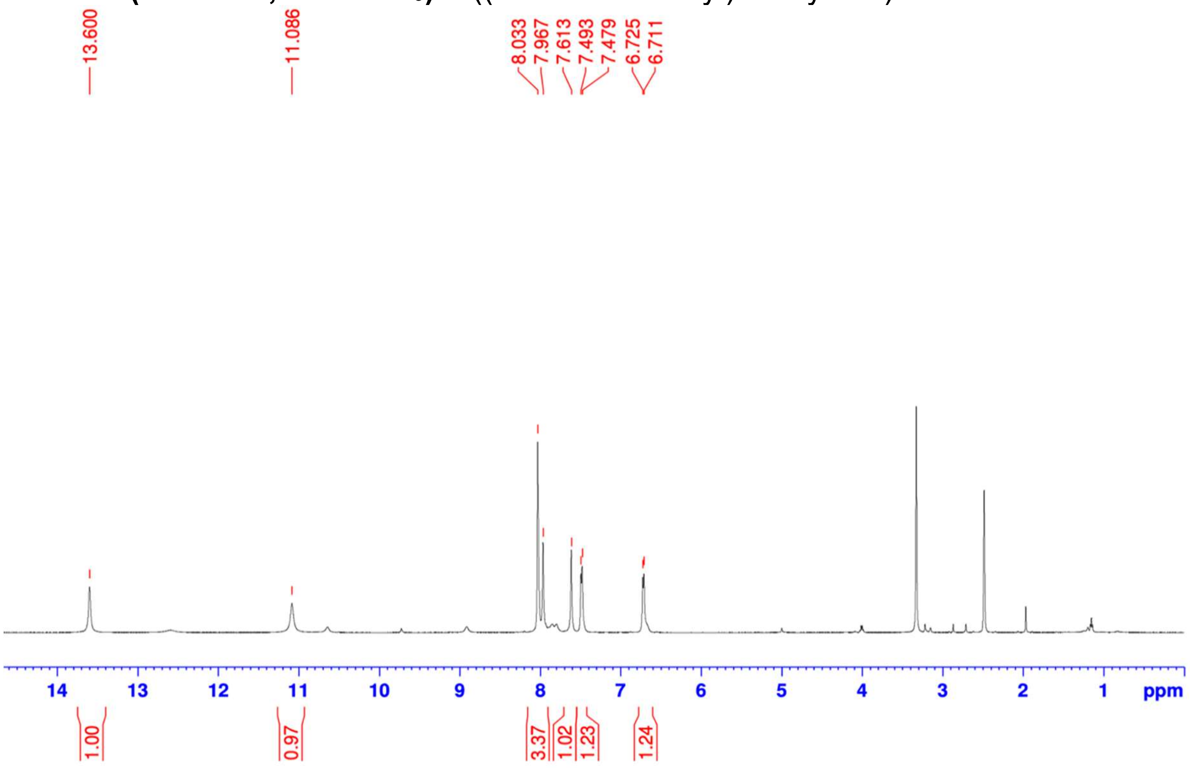


**$^{13}$C NMR (125 MHz, DMSO-*d₆*)** 3-((1H-imidazol-5-yl)methylene)-5-iodoindolin-2-one **2q**

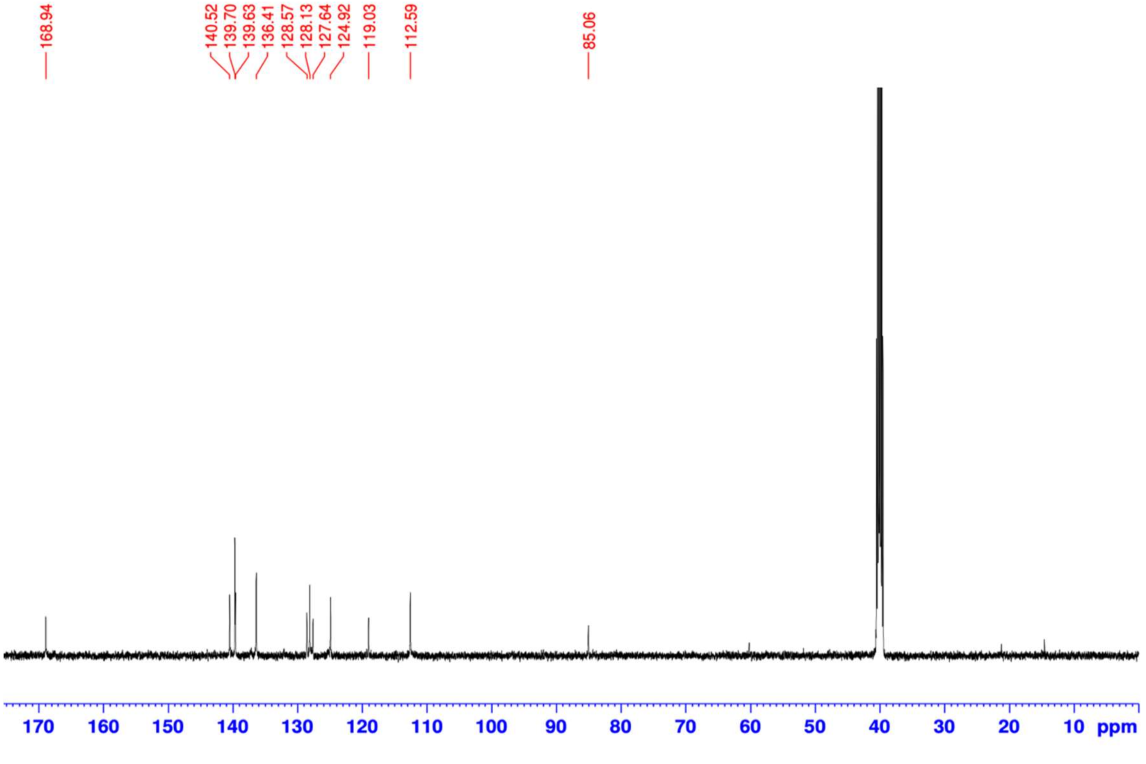

**$^1$H NMR (500 MHz, DMSO-$d_6$)** 3-(imidazo[1,2-a]pyridin-5-ylmethylene)-5-iodoindolin-2-one **2r**

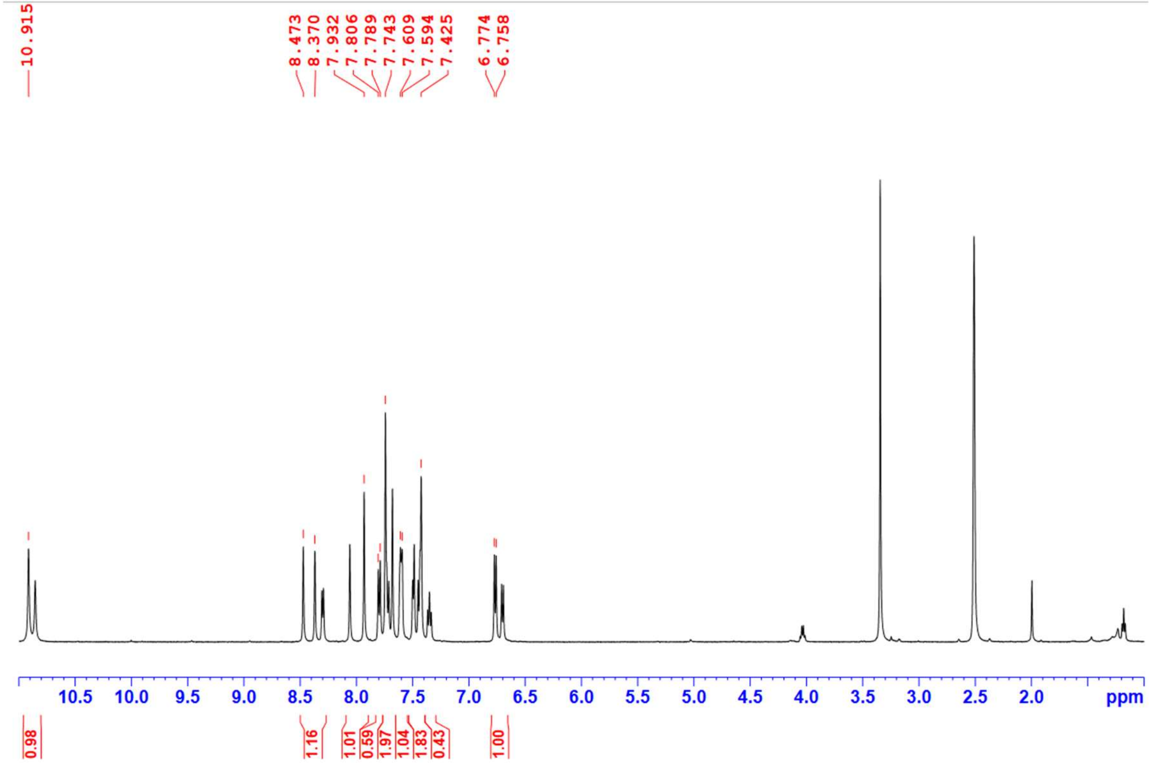


**$^{13}$C NMR (125 MHz, DMSO-$d_6$)** 3-(imidazo[1,2-a]pyridin-5-ylmethylene)-5-iodoindolin-2-one **2r**

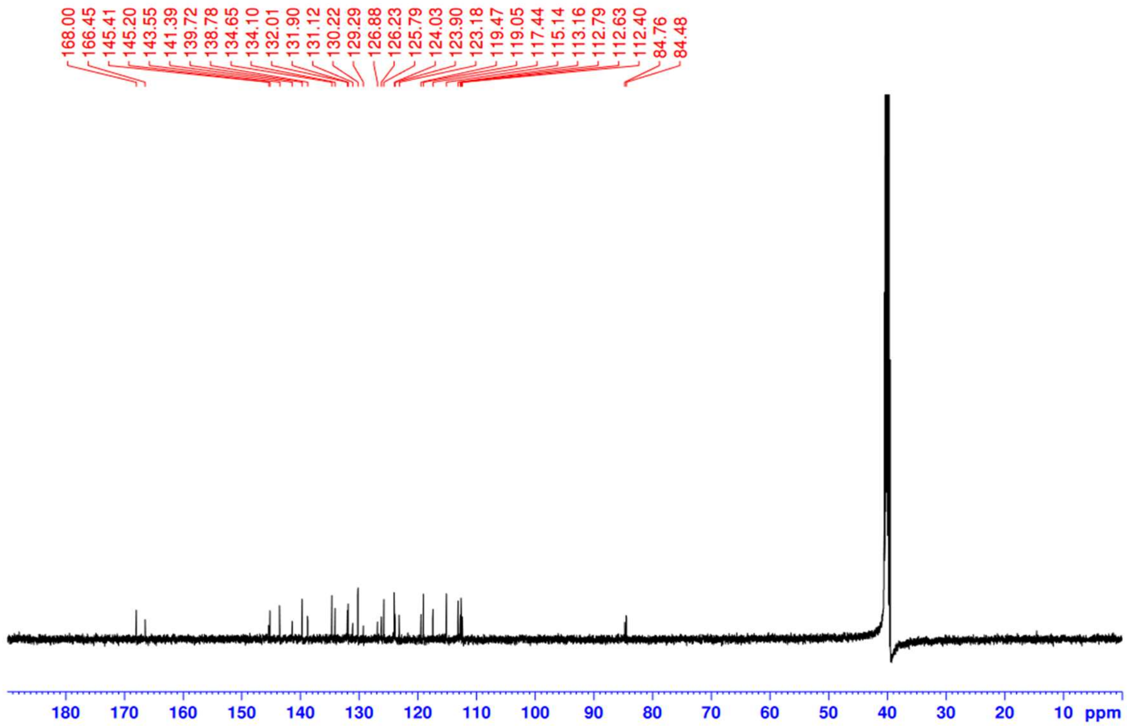

### 6. 2D-NMR Data

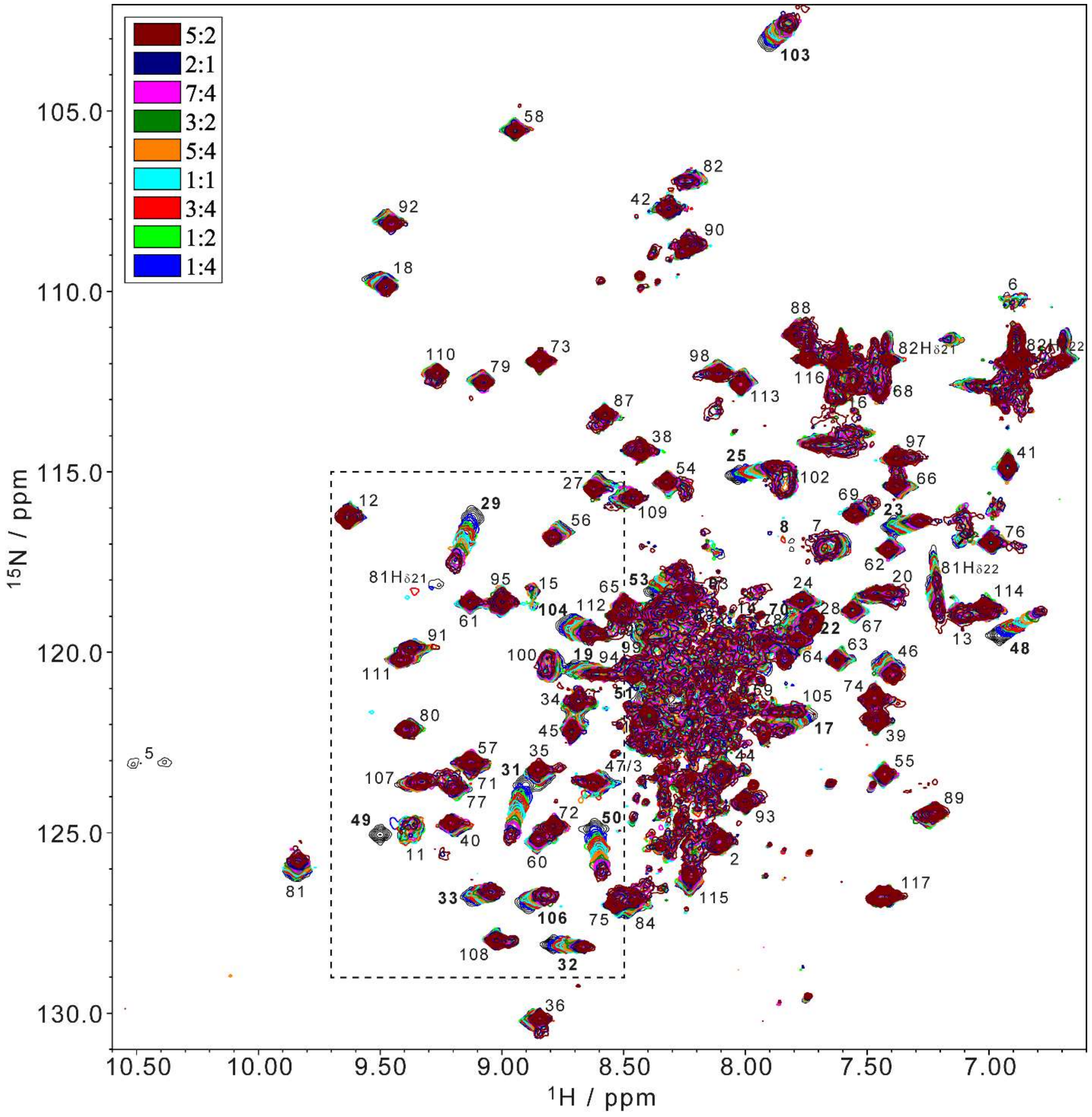


Overlay of the [$^{1}$H,$^{15}$N] HSQC spectra of a 200 μL sample of 200 μM [U-$^{15}$N] GABARAP with 20 mM sodium phosphate, 80 mM NaCl, 80 mM KCl, 40 μM EDTA in 10% (v/v) DMSO, 10% (v/v) $D_2O$, pH 6.9. Spectra were recorded at 25.0°C and 700 MHz alone (black) and in the presence of increasing amounts of a stock solution of compound **1** in the ratios indicated in the legend. Backbone resonance assignments of free GABARAP are indicated by residue numbers. Labels for amide side-chain resonances of Asn are shown as well, but those of Gln are omitted because of heavy overlap. Resonances exhibiting significant chemical shift perturbations upon ligand binding are labeled in boldface, and include E8, E17, E19, R22, K23, Y25, V29, V31, I32, V33, K48, Y49, L50, V51, S53, H70, F103, F104, and Y106. Crowded contour lines in the central region of the spectrum are due to the low contour lines selected for this plot, which were required to visualize clearly the resonances of GABARAP that are affected by line-broadening due to conformational dynamics such as Y5 and E8. Colors of the spectra match colors in Fig. 7, and the spectral region indicated by the dashed box is further shown in Figure 7b of the main text.
